# Heterogeneity in killing efficacy of individual effector CD8+ T cells against Plasmodium liver stages

**DOI:** 10.1101/2022.05.18.492520

**Authors:** Soumen Bera, Rogerio Amino, Ian A. Cockburn, Vitaly V. Ganusov

**Affiliations:** Department of Microbiology, University of Tennessee, Knoxville, TN 37996, USA; Unit of Malaria Infection and Immunity, Institut Pasteur, 75015, Paris, France; Division of Immunology, Inflammation and Infectious Disease, John Curtin School of Medical Research, The Australian National University, Canberra, 2600, Australia; Department of Mathematics, University of Tennessee, Knoxville, TN 37996, USA; Host-Pathogen Interactions program, Texas Biomedical Research Institute, San Antonio, TX 78258, USA

**Keywords:** CD8+ T cell, Plasmodium sporozoites, liver stages, intravital imaging, mathe-matical modeling

## Abstract

Vaccination strategies in mice inducing high numbers of memory CD8+ T cells specific to a single epitope are able to provide sterilizing protection against infection with Plasmodium sporozoites. We have recently found that Plasmodium-specific CD8+ T cells cluster around sporozoite-infected hepatocytes but whether such clusters are important in elimination of the parasite remains incompletely understood. Here we used our previously generated data in which we employed intravital microscopy to longitudinally image 32 GFP-expressing Plasmodium yoelii parasites in livers of mice that had received activated Plasmodium-specific CD8+ T cells after sporozoite infection. We found significant heterogeneity in the dynamics of the normalized GFP signal from the parasites (termed “vitality index” or VI) that was weakly correlated with the number of T cells near the parasite. We also found that a simple model assuming mass-action, additive killing by T cells well describes the VI dynamics for most parasites and predicts a highly variable killing efficacy by individual T cells. Given our estimated median per capita kill rate of *k* = 0.031*/*h we predict that a single T cell is typically incapable to kill a parasite within the 48 hour lifespan of the liver stage in mice. Stochastic simulations of T cell clustering and killing of the liver stage also suggested that 1) three or more T cells per infected hepatocyte are required to ensure sterilizing protection; 2) both variability in killing efficacy of individual T cells and resistance to killing by individual parasites may contribute to the observed variability in VI decline, and 3) the stable VI of some clustered parasites cannot be explained by measurement noise. Taken together, our analysis for the first time provides estimates of efficiency at which individual CD8+ T cells eliminate intracellular parasitic infection in vivo.

## 1 Introduction

Malaria is a mosquito-borne disease caused by parasites from genus Plasmodium; Plasmodium parasites of humans are responsible for more than 200 million infections and 500,000 deaths annually [1]. Infection starts when mosquitoes inject sporozoites (a specific stage of Plasmodium parasites) into dermal tissue [2, 3]. The sporozoites move through the tissue, find and invade blood vessels, and disseminate in the body via the blood. From the blood some sporozoites will invade the liver, infect hepatocytes and form liver stages. It takes approximately 2 days (in mice) or 6-7 days (in humans) for liver stages to develop into merozoites (another stage of malaria parasites) [4]. Merozoites infect red blood cells and ultimately cause clinical symptoms of malaria. Several vaccine candidates aim at preventing infection; including some that aim at removing all liver stages by inducing large numbers of Plasmodium-specific memory CD8+ T cells [5–7].

In an experimental setting multiple strategies have been used to generate large numbers of Plasmodium-specific CD8+ T cells including priming mice with dendritic cells (DCs) coated with Plasmodium peptides and boosting with Listeria monocytogenes (LM), expressing Plasmodium peptides, immunization with radiation-attenuated sporozoites (RAS), and adoptive transfer of in vitro activated CD8+ T cells [5, 8–13]. While these strategies have shown good protection following exposure to Plasmodium sporozoites (e.g., [5, 14]), specific mechanisms by which vaccine-induced CD8+ T cells find the infected cells and the critical parameters required for parasite’s killing such as the number and duration of CD8+ T cell contacts with parasitized cells, remain largely unknown.

Intravital microscopy has been used to gain quantitative insights into the process by which CD8 T cells kill their targets in vivo. Mempel *et al*. [15] directly observed the killing of targets (peptide-pulsed B cells) by CD8+ T cells in vivo. Interestingly, the authors found that formation of the conjugate between the T cell and its target eventually resulted in the arrest of target cell movement and death within 20-30 min of initial contact [15]. In contrast, the killing of cancer cells by adoptively transferred activated CD8+ T cells took on average 6 hours [16]. Subsequently we documented the kinetics of killing of individual parasitized cells by CD8+ T cells in vivo [17]. In that study we followed the kinetics of formation of clusters of Plasmodium-specific CD8+ T cells around Plasmodium liver stages in mice using intravital microscopy; interestingly, the formation of larger clusters (3 or more cells present within 40 *μ*m of the parasite) was associated with a higher chance of parasite death during the imaging period [17]. Also, based on the kinetics of the loss of fluorescence signal from the GFP-expressing parasites, we observed four different patterns of parasite’s death; however, the actual killing kinetics was not quantified [17]. Recent work extended our observations to CD8+ T cell-mediated killing of virus-infected cells suggesting that after initial contact, it takes on average 30 minutes for a T cell to kill its target [18]. Interestingly, while the initial analysis suggested that there may be cooperation between individual CD8+ T cells bound to the same virus-infected cells, further analysis suggested that such a conclusion may be an artifact of the data analysis that ignored prolonged “zombie” contacts of CD8+ T cells with already dying targets [18, 19]. Another study suggested that killing of cancer cells may require several contacts by the same or different CD8+ T cells occurring within a short (*<* 1 h) time period [20].

Following our previous work here we provide comprehensive quantitative analysis of the data on the kinetics of CD8+ T cell-mediated killing of Plasmodium liver stages. In these experiments we could accurately monitor the number of Plasmodium-specific CD8+ T cells located in the vicinity of GFP-expressing liver stages and “health” of the liver stages (indicated by a relative magnitude of the GFP signal) for several hours for 32 individual parasites [17]. Most of the liver stages (24/32) had at least a single CD8+ T cells nearby at some point of imaging, and our recently developed density-dependent recruitment model of T cell cluster formation could reasonably well describe the kinetics of accumulation of Plasmodium-specific CD8+ T cells near individual parasites over time. By fitting a simple mathematical model to the VI decline data we found that per capita killing efficiency of Plasmodium-specific CD8+ T cells in vivo varied dramatically between different parasites. Interestingly, we did not find any evidence that CD8+ T cells cooperate in parasite’s killing as the basic, mass action-based model could describe the data well for most parasites. Stochastic simulations suggested that given high variability in T cell killing efficacy, three or more T cells per liver stage are needed to ensure death of the parasite in 48 hours. Finally, we found that the variability in VI dynamics observed in the data suggests that many T cells around individual infected cells may be non-cytolytic (i.e., lack the ability to kill).

## 2 Materials and methods

### 2.1 Experimental Data

Experimental data for our analyses have been generated in our previously published work [17] except for the novel data on the dynamics of the vitality index (**VI**, see below) in control mice for a subset of parasites. In short, there were two sets of experiments performed (**Figure 1A**). In control experiments, Balb/c mice were infected with 3 *×* 10^5^ GFP-expressing Plasmodium yoelii (**Py**) sporozoites i.v., and 24h later the livers of mice were surgically exposed and imaged using a spinning disk confocal microscope. Imaging frequency varied between individual mice and/or parasites from 1.16 min to 5.07 min per z-stack. In total, 38 parasites in 2 different mice were followed for 6 hours. Imaging data from 22 of such parasites were analyzed longitudinally at multiple time points. Specifically, the vitality index (VI) of each parasite at different times was calculated in ImageJ as the log_10_ of the parasite brightness to the background brightness in the GFP channel using maximal z projection. In control experiments parasites were defined as superficial or deep based on their higher or lower z position in the stack of images, respectively (e.g., **Supplemental Figure S1**). In treatment experiments, mice were infected with 3 *×* 10^5^ GFP-expressing Py sporozoites. Twenty hours later, 9 *×* 10^6^ activated Py-specific CD8+ T cells were transferred into infected mice (for details of how activated Py-specific CD8+ T cells were generated see [17]). Four to eight hours later, livers of mice were surgically exposed and imaged using a spinning disk confocal microscope. In total, 32 parasites were followed in 4 mice. VI was calculated for each parasite as in control experiments, and by segmenting the movies with Imaris (https://imaris.oxinst.com), the number of T cells found within 40 *μ*m of the liver stage over time was quantified [21–23]. Following previous analyses we assume that the parasite is considered dead if the VI becomes less than 0.2 [17]. Our main results, however, were not particularly sensitive to this assumption (see Results section).

**Figure 1:**
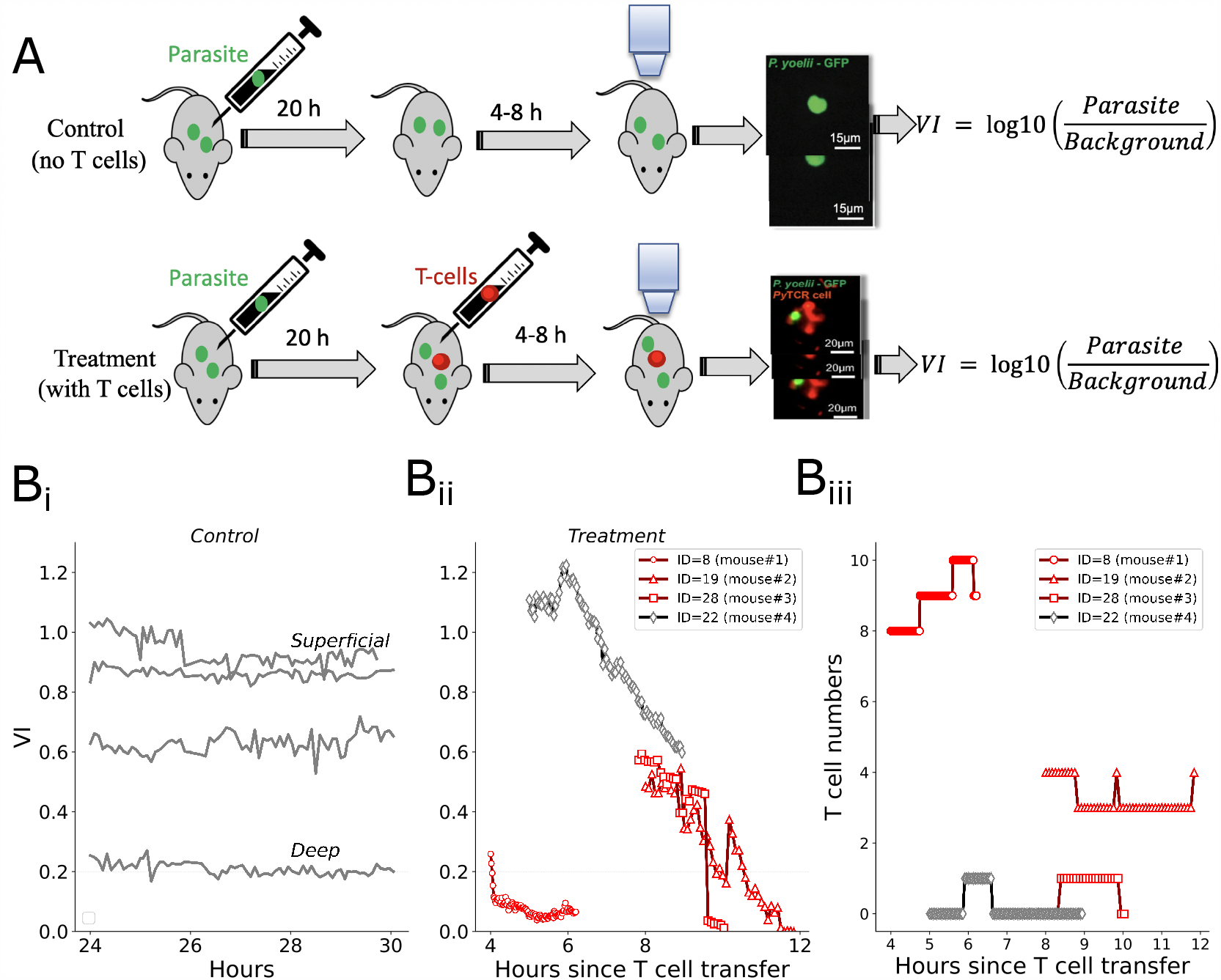
Experimental design to measure T cell impact on Plasmodium liver stages. **(A)** Experimental design. Mice were infected with 3 *×* 10^5^ GFP-expressing Py sporozoites and divided into two groups as control or treatment. In the control group mice were left until imaging started 24-28 hours after the infection. In the treatment group, 9 *×* 10^6^ Py-specific activated CD8+ T cells (**PyTCR**) were transferred into infected mice 20 hours after infection. Four to eight hours later, murine livers were surgically exposed and z-stacks of images were taken using spinning disk confocal microscope [17]. Vitality index (**VI**) of a given parasite was calculated as the log_10_ of the ratio between brightness of the GFP signal from the parasite to the brightness of the background based on maximal z projections. The number of T cells within 40 *μ*m of each liver stage was also calculated. **(Bi**,**Bii)**: Example of the VI dynamics for control (Bi) or treatment (Bii) groups. “Superficial” and “deep” denote location of the parasite based on the z coordinate in the imaging data. **(Biii)** Change in the number of Py-specific CD8+ T cells near the Py-infected hepatocytes over time since T cell transfer. See **Supplemental Figure S1** for all data graphs.

We provide all numeric experimental data as the supplement to the paper. These data include the following.

1. For control experiments, imaging was performed from 24 hour to 30 hour and from 30 hour to 33 hour after infection in 2 mice, respectively. The data include VI over time for each of 22 parasites (**Figure 1Bi** and **Supplemental Figure S1**A).
2. For treatment experiments, imaging was performed in different intervals that typically varied per mouse but also per parasite: 4 to 6.2 hours, 8 to 11.8 hours, 5 to 9.12 hours, 6 to 9.12 hours, 6.5 to 7.32 hours, and 7.82 to 11 hours in four different mice (**Figure 1**Bii-Biii and **Supplemental Figure S1**B-D). For every mouse and parasite ID, the data include the time since infection (hours), the number of T cells per parasite, GFP level of the background and the parasite, and the parasite’s VI.

In cases when VI or T cell number per parasite could not be calculated (due to imaging artifacts), values were indicated as NA. These data were excluded in the analysis.

### 2.2 Mathematical models

#### 2.2.1 Mixed effect modeling of VI dynamics

To separate the effect of time (*t*), T cell number per parasite (*T*), and mouse number (*M*) in the VI dynamics we performed linear mixed-effect modeling analysis. Using *lme4* package in R we considered the following alternative models that take into account time since T cell transfer *t* and the number of T cells per parasite *T* :

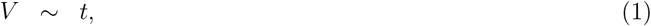

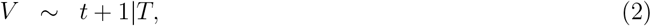

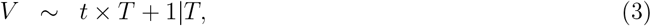

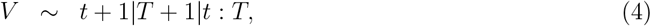

and to determine potential impact of individual mice on the VI dynamics we considered the following models:

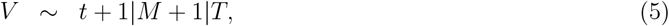

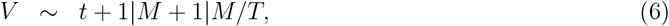

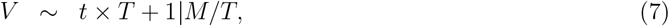

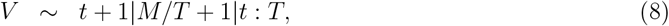

where *∼*represents regression, 1|*T* represents a random effect, 1|*M* is a random effect of four different mice, *t × T* is the main effect with interaction between time and T cell number, 1|*M/T* indicates a nested random effect of T cells from different mice, 1|*t* : *T* is a random effect with interaction between T cell number and time. This analysis was performed on 24 parasites that had at least one T cell within 40 *μ*m of the parasite at some time during imaging. All models were compared with the null model (no T cells or mouse effect) using ANOVA, and the best model was determined based on the lowest Akaike Information Criterion (AIC) value [24, 25].

#### 2.2.2 Mechanistic modeling of T cell clustering dynamics

We have recently shown that dynamics of CD8+ T cell cluster formation around the liver stages is best described by the density-dependent recruitment (**DDR**) model in which T cells enter the cluster at a rate *λ*_0_ + *λ*_1_*T* (*t*) where *T* (*t*) is the number of T cells near the parasite at time *t* and leave the cluster at a constant per capita rate *μ* [17, 23, 26]. By fitting the DDR model to the kinetics of formation of clusters around individual liver stages we found that the model in which recruitment rates *λ*_0_ and *λ*_1_ decline with time describes the data better than the model with constant parameters [23]. Based on these results we propose the following model with time-dependent rates of T cell clustering in the DDR model:

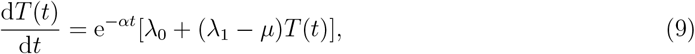

where *α* is the rate at which *λ*_0_, *λ*_1_, and *μ* decline with time since T cell transfer. Interestingly, this model fitted the data for all parasites with slightly lower quality as compared to the model in which recruitment-related parameters changed at 4 h post T cell transfer (AIC 182.1 vs. 177.7, Δ = 4.4, see Kelemen *et al*. [23] for details). The solution of eqn. (9) with the initial condition *T* (0) = 0 is

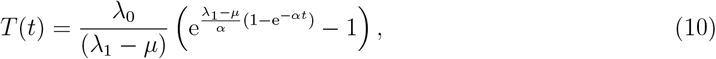

where only 3 parameter combinations (*λ*_0_, *λ*_1_ *− μ*, and *α*) determine the dynamics of the average cluster size. Unfortunately, experimentally measured cluster sizes for individual liver stages were insufficient to estimate all 3 parameters, in part because, faster recruitment rates *λ*_0_ and *λ*_1_ could be compensated by the faster decline in the recruitment rates given by *α*. Therefore, for further analyses we used a simplified DDR model in which we let *α →* 0:

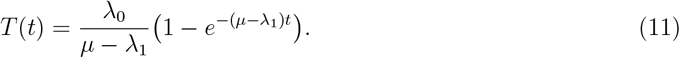

It should be noted that to describe typical cluster dynamics data with finite cluster size, condition *μ > λ*_1_ must be met, and only *λ*_0_ and the difference *μ − λ*_1_ can be estimated from the average cluster data. We found that predictions of the DDR model with constant and time-dependent parameters deviate insignificantly and thus, are not expected to influence estimate of the T cell killing rates. To estimate model parameters (*λ*_0_ and *μ − λ*_1_) we fitted eqn. (11) to the experimentally measured number of T cells per parasite over time for each parasite using least squares in Mathematica (using function NonlinearModelFit) or function nlsLM in R (using package minpack.lm) with control maxfev=10000 with maxiter=1024[27].

#### 2.2.3 Mechanistic modeling of VI dynamics

To further understand whether CD8+ T cells compete or cooperate in killing of the parasite and to estimate the per capita killing efficiency of Plasmodium-specific CD8+ T cells we used the following model for the dynamics of parasite’s VI

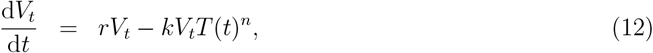

where *V*_*t*_ represents the parasite’s VI at time *t, r* is the rate of VI’s growth, *k* is the killing efficiency of CD8+ T cells and *n* is Hill coefficient, indicating deviation of the killing from mass-action. Parameter *n* allows to detect cooperativity (*n >* 1) or competition (*n <* 1) between different T cells in killing. We compare whether a simpler model (with *n* ≠ 1) describes the data better as compared to the full model (with *n* ≠ 1). The dynamics of the number of CD8+ T cells per parasite *T* (*t*) is given in eqn. (11). The models were fitted to VI data from individual parasites using least squares using the FindMinimum routine in Mathematica. For each model (two different functional responses with *n* = 1 and *n* ≠ 1) we performed the above procedure separately. We then used the AIC to select the best model from the two different functional CD8+ T cell responses:

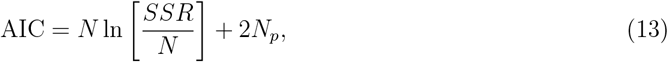

where *SSR* is the sum of squared residuals from the fit, *N*_*p*_ is the number of data points and *N*_*p*_ is the number of model parameters fitted to data [24, 25]. Confidence intervals for the best fit models were determined by resampling residuals with 1000 simulations [28].

#### 2.2.4 Stochastic model of killing of the liver stage assuming variability in T cell killing efficacy and/or parasite resistance

We simulated T cell-mediated killing of the liver stages by selecting a given number of T cells per parasite (parameter *n*_*T*_). Each of *n*_*T*_ T cells was assumed to either have maximal killing efficacy *k* = 0.1*/*h or to be nonlytic (*k* = 0) with probability *f*_*K*_. Parasites may be either fully sensitive to killing or fully resistant to killing by T cells; this was determined randomly for each liver stage with the probability *f*_*R*_. For each T cell cluster we allowed T cells to interact for *dt* h chosen uniformly between 0 and 1 h. We determined if the liver stage was eliminated by summing the killing efficacies of *n*_*T*_ T cells in the cluster and the time *dt* that T cells interacted with the liver stage assuming the killing process is exponentially distributed (per eqn. (12)). For each run we calculated the T cell killing rate as the ratio of the probability of liver stage death over the number of T cells and the duration of T cell interaction with the liver stage, 1*/*(*n*_*T*_ *dt*) or 0. We repeated simulations 10,000 times for the same number of T cells and calculated the average killing rate for a given set of model parameters.

#### 2.2.5 Stochastic model of variability of parasite’s VI over time

To investigate VI variability that may come from stochasticity of parasite’s dynamics we performed stochastic simulations of the parasite’s VI in the presence of different numbers of T cells near the parasite using Gillespie’s algorithm [29]. Simulations were performed to mimic experimental design. Specifically, in experiments the GFP signal from the parasite could theoretically vary between 0-255 in unit increments (8 bit), however, typically, by the start of imaging, the GFP signal rarely exceeded 200 with the background generally falling in the range of 10-20. For simplicity we assume that parasite’s health signal starts at 100 with the background being 1, resulting in the initial VI being log_10_(100*/*1) = 2. The parasite’s GFP signal was allowed to grow over time with a minimal growth rate (*r* = 0.0001*/*h) for twenty hours while the background remained fixed. At this time, T cells were transferred into mice and the T cells were allowed to accumulate at the parasite at the rate defined by parameters *λ*_0_ and *μ − λ*_1_ (see eqn. (11)) and killed the parasite at a rate *k*. The simulations were run for another 36 hours. Two hundred simulations were run using python package *gillespy2* with *Tau-Hybrid* solver. In another set of simulations instead of modeling increase in T cell numbers near the parasite we allowed T cell numbers to change randomly over time in the range observed for a given parasite.

## 3 Results

### 3.1 Experimental framework to quantify how *Plasmodium*-specific CD8+ T cells kill *Plasmodium yoelii* parasites in the liver

To understand how Plasmodium-specific CD8+ T cells may eliminate Plasmodium yoelii (**Py**) sporozoites from the liver we have performed intravital imaging experiments in which GFP-expressing parasites were imaged in the livers of live mice using intravital microscopy [17, **Figure 1**A]. From these imaging data, we calculated the relative brightness of the GFP signal from Py liver stages (denoted as vitality index or **VI**) in both control and treatment experiments (**Figure 1**Bi-Bii) and the number of Plasmodium-specific CD8+ T cells located within 40 *μ*m from the parasite at different times after T cell transfer in treatment experiments (**Figure 1**Biii).

In control experiments in which no T cells were transferred, VI remained constant for all except for one parasite that were followed longitudinally (**Supplemental Figure S1**A). Note that in our original analysis when parasite’s VI was measured only at 2 time points, VI for three parasites declined with time [17]. Estimated growth rates of the VI on average did not significantly differ from 0 (*p* = 0.92 and *p* = 0.13 for mouse 1 and 2, respectively, Sign-Ranked test) suggesting that in the absence of T cells, the VI does not increase over several hours of the imaging experiments. This is perhaps expected given that the VI denotes the relative GFP intensity and the parasite’s growth may be better estimated by the size of the liver stage [30]. Intriguingly, VI values differed among parasites, particularly those situated deeper in tissues (based on the slice’s z coordinate). Nonetheless, most values exceeded the previously suggested VI cutoff of 0.2, indicative of dead or dying parasites [17].

Importantly, the VI was more variable over time and between individual parasites in mice that had received Py-specific T cells (**Figure 1**Bii, **Supplemental Figure S1**B, **Supplemental Figure S2** and **Supplemental Figure S3**). In these experiments 8 out of 32 of the liver stages were not surveyed by T cells over the imaging period (i.e., no T cells were within 40 *μ*m of the parasite, **Supplemental Figure S1**C and **Supplemental Figure S4**). The change in the number of T in the vicinity of each parasite was variable with some parasites accumulating more T cells while others loosing T cells over time, and several parasites were only surveyed by a single T cell for a short period of time (**Figure 1**Biii and **Supplemental Figure S4**). Interestingly, many parasites had stable VI despite having multiple T cells nearby (e.g., **Supplemental Figure S3**). The regression analysis between the rate of VI decline and the average number of T cells per parasite showed a weak correlation (**Supplemental Figure S5**) suggesting that the impact of CD8+ T cells on the VI dynamics is surprisingly weak and highly variable between individual parasites and mice.

### 3.2 The number of T cells per parasite negatively correlates with parasite’s VI

To better understand how the number of CD8+ T cells per parasite is related to the parasite’s VI and how imaging of individual mice may influence the results, we performed several alternative analyses. First, we analyzed the relationship between the VI and the number of T cells per parasite for each mouse. Interestingly, we found that a higher number of T cells per parasite was negatively correlated with the parasite’s VI for three out of four mice (**Figure 2**A). There was no correlation between CD8+ T cell numbers per parasite and parasite’s VI for many parasite-T-cell combinations, in part, because for many parasites the number of T cells did not vary over time or there were no T cells near the parasite (13/32). For the 19 parasites, most correlations (12/19) between VI and CD8+ T cell number were negative (**Figure 2**B) suggesting that at least for some parasites increasing numbers of CD8+ T cells in the cluster reduce parasite’s “health”. Because for some parasites the number of T cells did not change over time we re-analyzed the data using a novel concept of dynamical correlations [31] by using `dynCorr` library in R. Interestingly, the overall correlation for all 32 parasites between VI and T cell number was negative (-0.38). However, the bootstrap analysis (function `bootstrapCI` in `dynCorr`) indicated that the correlation is not statistically different from 0 (estimated 95% Cis = -0.55 - 0.56). We found similar results when restricting the analysis to 19 parasites for which the number of T cells per parasite varied over time.

**Figure 2:**
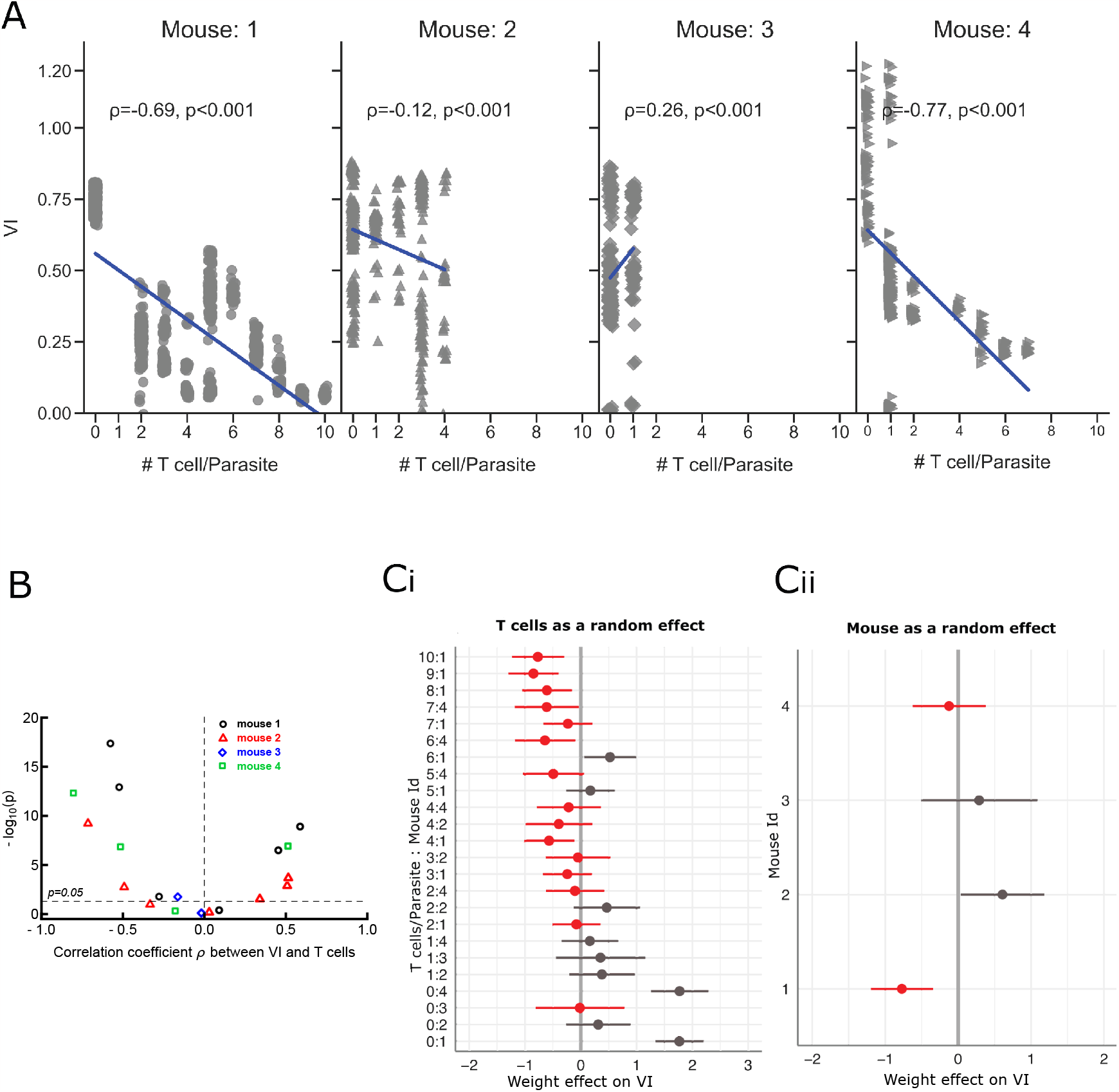
Higher CD8+ T cell numbers per parasite are negatively correlated with the parasite’s VI. (**A)** Correlation between parasite’s VI and T cell numbers per parasite is shown for individual mice. The Spearman rank correlation coefficients *ρ* and corresponding p value were calculated using *scipy*.*stat* package in python and are shown on individual panels. (**B**) Spearman rank correlation between parasite’s VI and CD8+ T cell numbers per parasite is shown for all parasites for which such correlation existed along with estimated p value from the correlation test (19/32). Dashed horizontal line denotes *p* = 0.05. (**Ci**) Prediction of the best fit linear mixed effects model assuming T cell numbers per parasite as a random effect (eqn. (6)) are shown for individual T cell number:mouse combinations. Negative values for effect (weight) correspond to T cell numbers that reduce VI over time. (**Cii**) Prediction of the best fit linear mixed effects model assuming mouse as a random effect (eqn. (5)) are shown for individual mice (negative and positive effect corresponds to the parasite’s VI declining or increasing, respectively). In panels C, grey and red colors denote positive and negative weight values, respectively.

To understand how the time since T cell transfer may impact the relationship between T cell number per parasite and parasite VI we fitted several alternative models to the parasite VI normalized to its average value for each parasite for all mice but dividing the data into different time periods (see **Figure 1**Ciii, **Supplemental Figure S6**, and eqns. (S.1)–(S.5) in Materials and methods). Specifically, we divided the VI and T cell data for the 32 parasites into four different cohorts based on the time interval since T cell transfer (4-6 hours, 6-8 hours, 8-11 hours, and all data combined, 4-11 hours) and fitted five alternative models (eqns. (S.1)–(S.5)) to these data using least squares. All models described the data with similar quality and model predictions were typically similar for the alternative models with the exception of the exponential model that gave slightly different predictions (**Supplemental Figure S6**A-C). For each time period we then calculated the number of T cells, *T*_*IC*50_, that was needed to reduce parasite’s VI to 50% of its initial value. Interestingly, *T*_*IC*50_ increased with the time since T cell transfer from about 1 T cell per parasite at early time points (4-6 hours) to 13 T cells per parasite at later time points (8-11 hours, **Supplemental Figure S6**A-C). This result may indicate that there may be something special about parasites that had survived for 8-11 hours of T cell search for infection; perhaps such parasites are able to actively resist killing or they are located in areas of the liver that are less accessible to T cells. When analyzing all the data together we found that about 3 Py-specific CD8+ T cells per parasite were needed to reduce the parasite VI to 50% of its initial value (**Supplemental Figure S6**D).

Finally, to quantify the effect of the time since T cell transfer, the number of T cells per parasite, and individual mouse on the VI dynamics we performed several linear mixed model-based analyses (see eqns. (1)–(8) and Materials and methods for more detail). By comparing among alternative models, the model in which VI was dependent on time since T cell transfer as a fixed effect and mouse being a random effect (eqn. (6)) described the data with best quality as perhaps one would expect. In this model, based on total variance explained by the model, the overall contribution of CD8+ T cell number on parasite’s VI was about 60% and the mouse effect was around 40%. Overall, higher T cell numbers were associated with a decline in the observed VI over time (**Figure 2**Ci); however, this result was mouse-dependent (**Figure 2**Cii). This analysis suggests that the number of CD8+ T cells around individual parasites only partially explains the dynamics of the parasite VI, and other factors – for example, killing of the parasite prior to imaging or location of the parasite in a deeper tissue – may be also important.

### 3.3 A simple model with mass-action killing describes well the VI dynamics for most parasites

Having performed several statistical analyses of the data we next turned to a more mechanistic modeling of T cell-mediated killing of parasites by asking whether the kinetics of VI loss over time can be accurately described by different mathematical models. We focused this analysis on the subset of parasites for which there was at least one T cell present near the parasite during the imaging time period (24/32 parasites); this excluded 8 parasites with no T cells from the analysis. Because imaging modalities only allowed us to observe T cell clusters at later points after T cell transfer and some T cells in the cluster may have initiated parasite killing before the imaging started, we needed a mathematical approximation to describe kinetics of T cell cluster formation after T cell transfer but prior to imaging start.

We have previously shown that a “density-dependent recruitment” (**DDR**) model best describes the clustering of CD8+ T cells around Plasmodium liver stages [17, 23]; this model proposes that recruitment of T cells to the liver stage increases with the number of T cells already present near the liver stage. Analysis of kinetics of cluster formation for liver stages in the whole dataset suggested that parameters of the DDR model are likely to change with time indicating reduced rate of T cell recruitment at later times [23, eqn. (9)]. However, fitting the DDR model with time-dependent parameters (DDR-alpha model) to data for T cell clustering around individual liver stages typically resulted in overfitting and did not allow us to estimate all 3 model parameter combinations (results not shown). Therefore, we fitted the version of the DDR model with constant parameters (eqn. (11)) to the measured numbers of CD8+ T cells per parasite using nonlinear least squares and estimated model parameters (*λ*_0_ and *μ − λ*_1_). The model fitted the data well (**Supplemental Figure S7**) and predicted relatively rapid formation of T cells clusters for most parasites as expected from our previous analysis [23]. Importantly, both estimated parameters varied dramatically between individual parasites (shown in a supplemental table available as a csv file) with median rates *λ*_0_ = 0.40*/*h and *μ − λ*_1_ = 0.13*/*h. By fitting the DDR model to the T cell clustering data for all parasites using nonlinear mixed effect approach we found that the data are best explained by a random effect of the constant recruitment rate *λ*_0_ (median *λ*_0_ = 0.33*/*h) and a fixed effect of cluster amplification rate *μ − λ*_1_ = 0.29*/*h (results not shown).

We then compared how well alternative simple models that use the estimated dynamics of T cell numbers per parasite but assume different killing terms, fit the data on the dynamics of VI (**Figure 3**A). Specifically, we fitted the models to the change in VI over time using nonlinear least squares. These alternative models assumed 1) mass-action killing rate (eqn. (12) with *n* = 1), 2) powerlaw killing term (eqn. (12)) that allows to detect cooperativity (*n >* 1) or competition (*n <* 1) between individual T cells in the cluster. In the model fits we assumed that the parasite’s growth rate *r* = 0 since our analysis of the data on VI dynamics in control mice found that this growth rate on average was not different from zero (**Supplemental Figure S1**A). In addition, we found that allowing *r* to be fitted resulted in overfitting of the model to VI data (results not shown). Interestingly, we found that the model assuming mass-action killing described the data with sufficient quality for the majority of the liver stages (**Figure 3**B-F and **Supplemental Figure S8**). Some liver stages did not display any VI decline despite being surrounded by T cells (e.g., **Supplemental Figure S8**J, V, X) but most parasites did display some loss of VI suggesting that our mathematical modeling-based approach allowed us to detect weak killing signals in these data. Importantly, we found dramatically variable per capita T cell killing rates ranging from *k* = 0*/*h to *k* = 2*/*h with the median *k*_med_ = 0.031*/*h (**Figure 3**G). Importantly, the estimated per capita killing rate did not depend on the average number of T cells observed for each liver stage for the imaging period (*ρ* = 0.03, *p* = 0.89) with some variability in median killing rates between individual mice (**Figure 3**G). The distribution of estimated killing rates is skewed with T cells around a small number of parasites showing very high kill rates while T cells around most parasites have low killing efficacy (**Figure 3**G).

**Figure 3:**
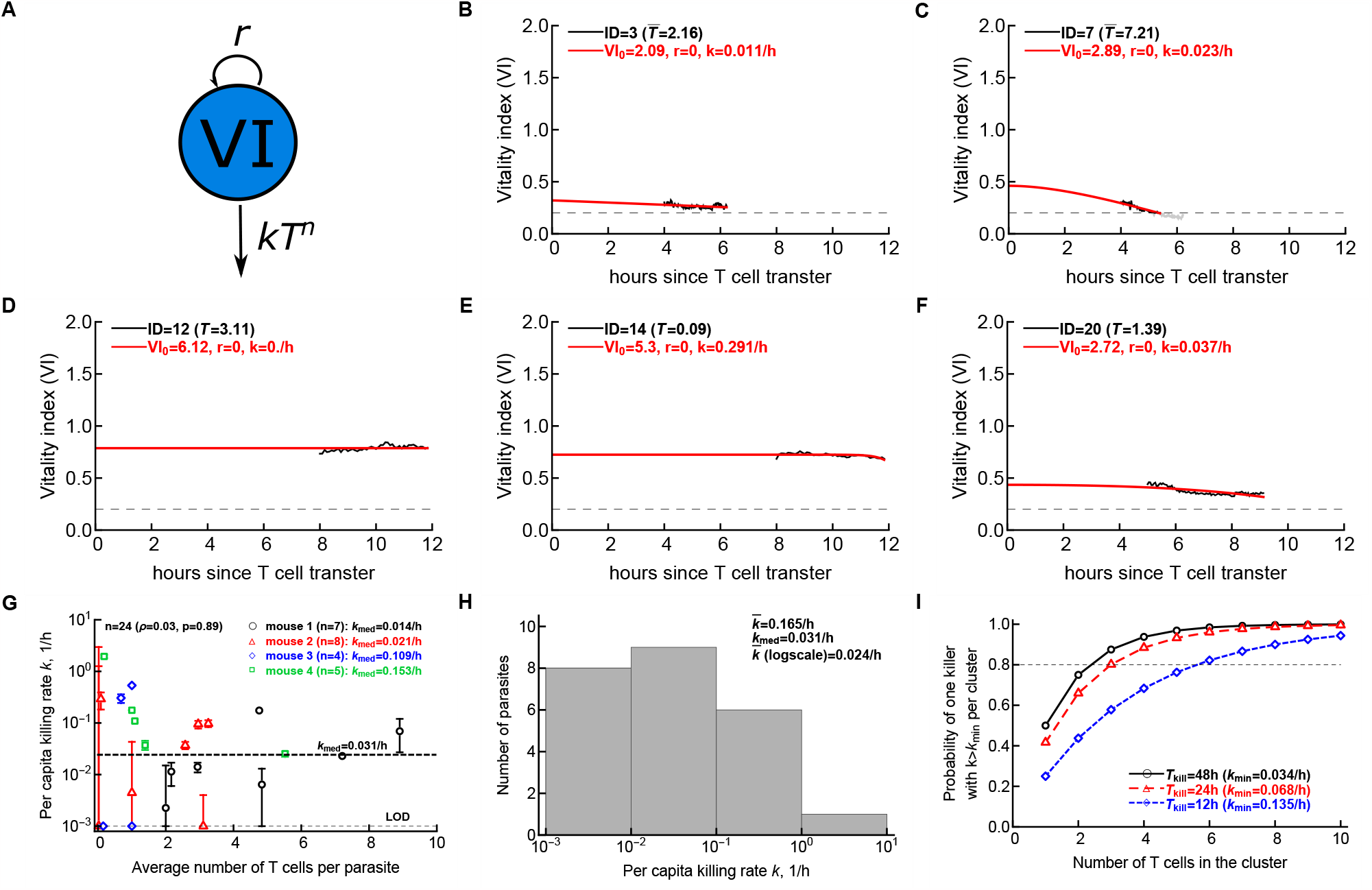
The best fit model suggests a relatively low killing efficacy of CD8+ T cells in vivo. (A) Schematic of the model depicting the essential interaction between CD8+ T cells (*T*) and VI (*V*_*t*_) where *k, r*, and *n* correspond to killing efficiency per T cell, growth rate of the parasites, and parameter indicating competition (*n <* 1) or cooperation (*n >* 1) between T cells located near the same parasite, respectively. Two alternative models (eqns. (12)–(12)) were fitted to the data on VI dynamics while taking into account change in the number of CD8+ T cells per parasite as predicted by the density-dependent recruitment model (**Supplemental Figure S7**). (B-F) Experimental data on VI change in subset of parasites are shown by markers and lines are the best fit predictions of the simplest, mass-action model (eqn. (12)). Estimated killing rate *k* is shown on individual panels, and all fits are shown in **Supplemental Figure S8**. The dotted horizontal line denotes the death threshold VI = 0.2. (G) Estimates of per capita T cell killing efficacy along with predicted 95% confidence intervals is plotted against the time-average of the number of T cells near each liver stage for *n* = 24 parasites which had at least one T cell at some time during imaging. Estimates with *k* = 0 were assigned value of 10^*−*3^*/*h (limit of detection, **LOD**) to display on the log-scale. Median estimates for the killing rate for individual mice or all data together (dashed line) are indicated on the panel. (H) Distribution of the estimated killing rates from G shown as a histogram. (I) Stochastic simulations on resampling of estimated killing efficacy per T cell from H predicting the probability of liver stage elimination within different time intervals (48h, 24h, or 12h) as the function of the number of T cells in the cluster. Horizontal dashed line denotes 80% probability.

To confirm the low median killing rate of CD8+ T cells we also fitted the mathematical model (eqn. (12) with *n* = 1) to the VI dynamics using nonlinear mixed effect approach in R (using `nlme` routine). In this analysis we used average number of T cells per parasite for each of 24 parasites (and not their dynamics) and allowed for fixed and random effects for the T cell killing efficacy and initial VI. Interestingly, we found the best model when indeed both *k* and initial VI were random effects. The best fit median killing rate was *k*_med_ = 0.035*/*h that was consistent with the estimate found by fitting the models to the VI dynamics of individual parasites.

The kill rate of *k*_med_ = 0.031*/*h by a “median” T cell suggests that it would take nearly 52 hours for such a cell to reduce parasite’s VI from 100% to 20% and thus, to kill the parasite (*T*_kill_ = 1*/k*_med_ ln(100*/*20) = 51.8h). Since this time is longer than the typical time of a liver stage (45-48h, [32]) this result suggests that a single CD8+ T cell is typically insufficient to eliminate a single liver stage. To understand how many T cells would be needed to eliminate an individual liver stage we resampled the estimated killing rates from **Figure 3**H for different number of T cells per parasite and estimated if for additive (mass-action) killing by *n*_*T*_ T cells, the parasite will die within 12, 24, or 48 h (**Figure 3**I). As expected, having more T cells in the cluster increases the chance of parasite’s death. Importantly, even for the longest possible time of contact between T cells in the cluster and the parasite (48h) three T cells were required to ensure the death of the liver stage with over 80% probability (**Figure 3**G). These results strongly suggest that while a single CD8+ T cell is clearly capable of eliminating a liver stage parasite (e.g., **Supplemental Figure S8**Q), clusters involving 3 or more T cells are needed to ensure elimination within 48 hours.

We also fitted an alternative model (eqn. (12) with *n ≠* 1) to the VI data and found that for several parasites the alternative model provided somewhat better fits as judged by the change in AIC values (IDs = 7, 14, 18, 20, 23). Visually this improvement in the fit allowed the model to more accurately describe exponential decline in VI with time for this subset of the liver stages (results not shown). Interestingly, in all cases the best fit resulted in *n <* 1 (typically *n* ≈ 0) suggesting competition between individual cells and saturation in killing after one T cell located the parasite. By fitting eqn. (12) to the VI dynamics data using nonlinear mixed effects approach improved the fit non-significantly while predicting *n* = 0.75. Thus, for most parasites the model assuming additive, mass-action-like killing was sufficient to accurately describe the data.

### 3.4 Stochastic simulations suggest that both variability in T cell killing efficacy and liver stage resistance to killing may contribute to the variability in estimated killing rates

The relationship between estimated per capita killing rate *k* and the average number of T cells observed during imaging for each liver stage (**Figure 3**G) had at least two interesting features: 1) the estimated killing rate varied orders of magnitude between T cells located near different liver stages, and 2) variability in the estimated killing rate is lower when larger numbers of T cells are in the cluster. To investigate the potential causes of these two features we ran stochastic simulations (**Supplemental Figure S9**A). In the simulations we allowed *n*_*T*_ T cells to interact with the liver stage that may be fully susceptible or fully resistant to killing by T cells (with a probability *f*_*R*_). T cells also may either be efficient killers (with *k* = 1*/*h) or be nonlytic (*k* = 0) sampled with a probability *f*_*K*_. We then allowed clusters of T cells to interact with the parasite for uniformly chosen lengths of time *dt* and calculated whether such interactions resulted in the parasite’s death. Repeating the simulations for 10^4^ liver stages for each parameter combination we could qualitatively recapitulate 1) variability in the estimated killing rate, and 2) decline in variability (given by the width of confidence intervals) with increasing cluster size (**Supplemental Figure S9**B-D). These simulations suggest that both the variable T cell killing efficacy of individual cells and the resistance of the liver stage to killing may contribute to the observed heterogeneity in the VI dynamics in vivo. Using mice allowing to detect events of T cell recognition of an infected cell (e.g., Ca-flux mice, [20]) may help discriminate between these alternatives.

### 3.5 Stochastic model with independent and random T cell response is unable to explain the stability of VI most parasites

In our set of 32 parasites several parasites displayed no evidence of decline in VI despite being surrounded by one or several CD8+ T cells for a period of time (e.g., parasite ID 11 and 12 in **Supplemental Figure S2**). Given that the estimated killing rate of CD8+ T cells was relatively low (**Figure 3**G) we hypothesized if constant VI could be explained simply by insufficient imaging period of a few hours preventing us to see a noticeable decline in the VI.

To test this hypothesis we performed stochastic simulations of the parasite’s GFP signal that would mimic our experimental design (see Materials and methods for detail). We started the simulation with the parasite’s GFP signal being 100 that could change up or down by one (similar to the information provided in the imaging data) and then calculated parasite’s VI assuming background of one (i.e., the initial VI was log_10_(100*/*1) = 2). We allowed the parasite VI to grow for 20 hours with a small growth rate *r* = 0.0001*/*h. After 20 hours T cells were transferred into the mice and started forming clusters around the parasite – the number of T cells per parasite was given by eqn. (11) with parameters chosen for individual parasites (**Supplemental Figure S7**). We started recording the VI 4-8 hours after T cell transfer and continued for another 2-4 hours. T cells were assumed to reduce the VI according to the mass action model with *k* = 0.031*/*h. In this way we simulated 250 trajectories and estimated the rate at which VI was declining with time for individual runs taking the time period of imaging into account (**Figure 4**). The VI for individual runs typically declined with time whether few or many T cells were located near the parasite (**Figure 4**A&C), though there was large variability in the decline rates (**Figure 4**B&D). Importantly, none of the slopes of VI decline in simulations were close to the slow VI decline rate observed in the data for “non-dying” parasites. We also performed stochastic simulations assuming that the number of T cells per parasite is random at different time point sampled from the range of T cell numbers observed for a given parasite. These simulations still failed to capture the declining slope of VI observed experimentally (**Supplemental Figure S10**). This suggests that our hypothesis that stable VI is due to stochasticity in the VI dynamics is not consistent with the data. The reasons why some parasites remain viable with no evidence of declining VI despite being surrounded by T cells remain unclear but may include at least three alternative explanations: 1) while being close, Py-specific CD8+ T cells do not recognize the infection, 2) the parasite was actively inhibiting killing signals that CD8+ T cells may be delivering to the infected cell, and 3) CD8+ T cells in the cluster are nonlytic. While we favor hypothesis #3, future experiments will be needed to rigorously discriminate between these alternatives.

**Figure 4:**
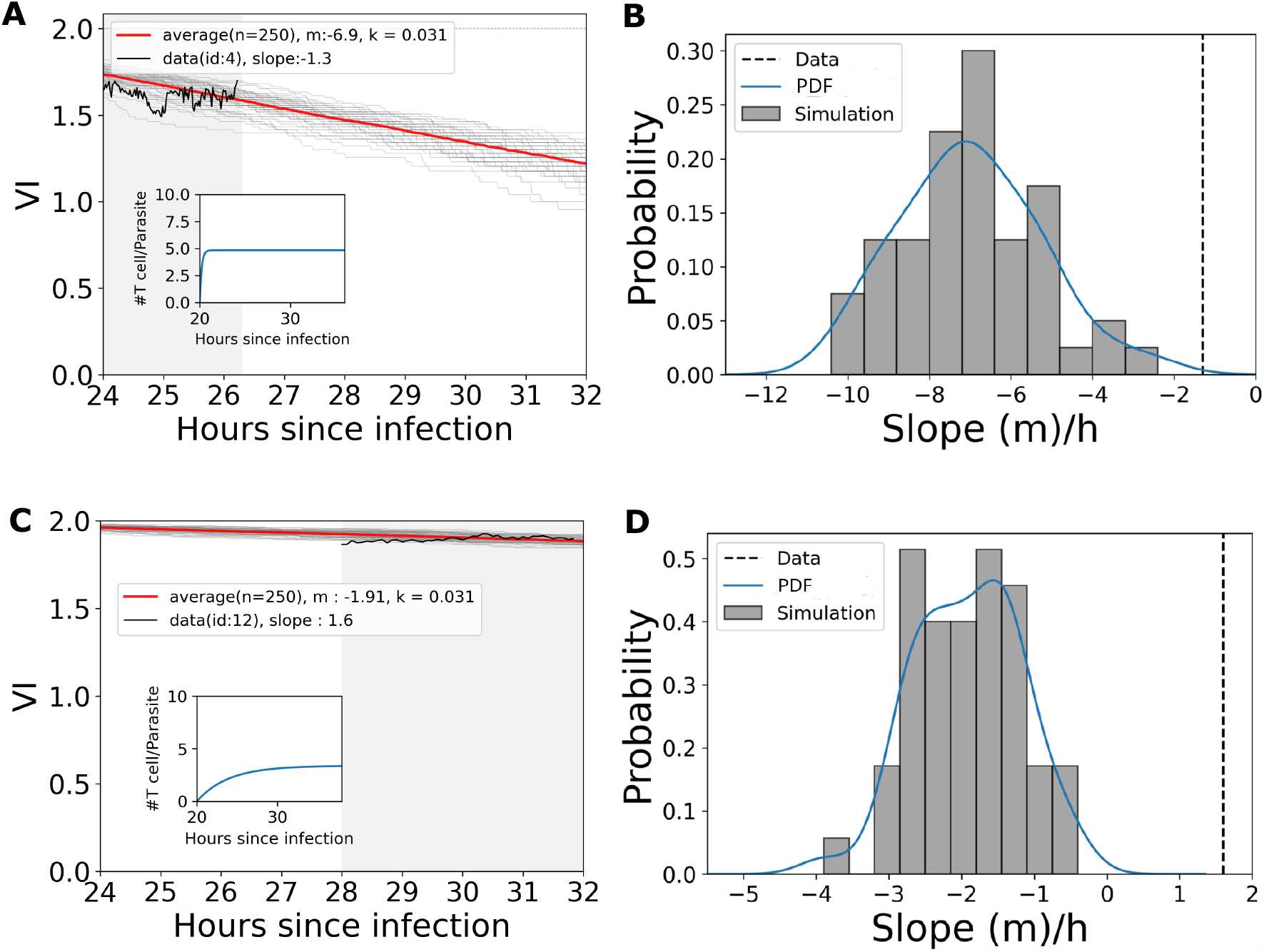
Constancy of parasite’s VI is not consistent with stochastic simulations of killing by T cells. We simulated parasite’s VI dynamics using Gillespie algorithm for 20 hours with a small growth rate *r* = 0.0001*/*h and then added killing by T cells at a per capita rate *k* = 0.031*/*h with the total kill rate being determined by the number of T cells per parasite increasing with time (see Materials and methods for more detail). **A, C**: We plot trajectory from 250 simulations (gray lines) with the average change (red line) calculated from every time step of 0.02 hour. T cell dynamics was added in the simulations using eqn. (11) with parameters estimated for individual parasites (A-B: ID = 4, C-D: ID = 12). Black lines in panels A&C show the actual change in VI over time as was observed in the data for parasite IDs 4 (A) and 12 (C). **B, D**: For every simulation we calculate the decay rate of the VI and show the distribution of decay rate with probability density function (blue line). The actual decay rate for the VI in the data for these two parasites is shown by horizontal black dotted line. Inserts show the T cell dynamics as assumed by the clustering model. Gray shaded area denote the imaging period which we used to calculate the VI decay rate. Simulations were done using package *gillespy2* with *Tau-Hybrid solver* in python. None of VI slopes in simulations were above the VI slope in the data suggesting *p <* 0.01 for the comparing simulations with the data.

## 4 Discussion

It is well established that large numbers of Plasmodium-specific CD8+ T cells are able to provide sterilizing protection against exposure to Plasmodium sporozoites but the mechanisms by which these T cells eliminate the parasites in the liver remain unclear. We used data from our previous intravital imaging of livers in live mice to accurately quantify kinetics at which Py-specific CD8+ T cells eliminate Py parasites from the liver [17]. In several alternative statistical analyses we found that the presence of Py-specific CD8+ T cells typically results in a faster loss of the VI signal from the parasites which we interpret as the process of parasite’s death. We estimated that on average 3 T cells per parasite are needed to reduce the parasite’s VI to 50% of its initial value during the imaging period (*∼*4 h), and that the dynamics of VI is better explained by the model in which the number of T cells per parasite steadily increases over time (**Supplemental Figure S6** and **Figure S11**). Analysis of the kinetics of T cell clustering and associated VI dynamics suggested that for most parasites T cells kill the parasite additively, via mass-action at a relatively low per capita killing rate of *k* = 0.031*/*h (**Figure 3**G) suggesting that a single CD8+ T cell will need around 52h to eliminate a liver stage. Thus, a single CD8+ T cell per liver stage is unlikely to be sufficient for sterilizing protection. There was large heterogeneity in the kinetics of VI decline between individual parasites; as such the rate of VI decline only weakly correlated with the number of T cells present near the parasite (**Supplemental Figure S5**). The VI of some parasites surrounded by several CD8+ T cells remained relatively constant during the imaging suggesting extremely small killing rate by the clustered T cells (**Figure 3**H). In agreement with this, stochastic simulations suggest that estimated variability in T cell killing efficacy require 3 or more T cells to cluster around a liver stage to ensure parasite elimination over the life span of the parasite (**Figure 3**H). Additional stochastic simulations of VI dynamics in presence of different numbers of T cells suggested that constancy of the VI over the observation period for some parasites cannot be due to limited duration of imaging (**Figure 4** and **Supplemental Figure S10**) and thus may be due to a lack of cytotoxic T cells in the cluster.

Details of how individual CD8+ T cells kill their targets in vivo remain poorly understood, in part of difficulty of recording killing events in live animals. Several studies reported that CD8+ T cells are able to kill either peptide-pulsed or virus-infected cells in 1-2 hours after initial contact [15, 18]. It may take longer times, about 6 hours, for a CD8+ T cell to kill a cancer cell [16] although a recent study suggests a larger kill rate of B16 cancer cells by CD8+ T cells in vivo [20]. Yet, all these estimates of kill rates are faster than found in our study suggesting that the type of target cell (e.g., type of intracellular pathogen) and of the tissue may influence the killing ability of T cells. One previous study found evidence of T cell cooperativity in killing of virus-infected cells [18], although a later re-analysis of the same imaging data suggested additive (mass-action like) killing by T cells [19]. Linear/additive effects of killing were also proposed in another study where delivery of 3 independent lethal hits by the same or different T cells within a short time period (*<* 1 h) was associated with target cell death [20]. We also did not find any evidence of T cell cooperation at elimination of Plasmodium liver stages but when multiple T cells cluster around the parasite, the probability of the parasite’s death during the imaging period was higher (**Figure 2**Ci). in contract, a recent study followed killing of tumor cells in individual organoids co-incubated with activated CD8+ T cells and found evidence of cooperativity of individual T cells at killing the tumors [33]. Whether this difference is due to different types of targets (infected hepatocytes vs. tumors) remains to be investigated.

Many studies have estimated the rate at which a population of antigen-specific CD8+ T cells kill their targets in vivo, for example, by using in vivo cytotoxicity assays (e.g., [34–37]), with target cell death rates ranging from minutes to days (reviewed in [38]). The exact reasons for this variability have not been fully elucidated but may involve differences in the type of target cells used in experiments, how T cell antigens are expressed by the targets (e.g., peptide-pulsed vs infected), and variability between individual T cells in their killing ability. Indeed, several recent reports tracking individual CD8+ T cells suggested great variability among antigen-specific T cells in their ability to kill their targets in vivo [18, 20]; moreover, variability in ability of different T cells to kill targets in vitro has also been well documented [20, 39, 40].

Our study has several limitations. To track the viability of the Plasmodium liver stages we imaged a relatively small area around the parasite (*∼*200 *×* 200 *×* 50 *μ*m) and T cells within 40 *μ*m of the parasite were considered to be close to the infected hepatocyte for killing [23]. For the parasites located at the deeper liver areas (with a higher z-coordinate), we may have underestimated the number of clustered T cells. Indeed, artificial increases in the number of clustered T cells improved the model prediction of VI dynamics supporting the idea that there may have been more clustered T cells than that observed experimentally. To more accurately evaluate how Plasmodium-specific CD8+ T cells eliminate the infection our experiments were performed in “reverse”: mice were first infected, and activated T cells transferred only 20 hours after the infection and with imaging performed several hours after the transfer of the T cells (**Figure 1**A). In part, this was also done due to the low GFP expression by individual Py sporozoites that would preclude imaging of individual parasites at the start of infection. This setup corresponds better to cases of adoptive T cell transfer therapy against cancer rather than prophylactic vaccination when activated/memory T cells are typically present prior to infection. Whether killing of Plasmodium liver stages proceeds similarly in cases when T cells are present in mice prior to infection and when sporozoites just invade the liver remains to be investigated.

We found that the estimated killing efficacy of T cells varied not only for individual parasites but also between mice; specifically, mouse 3 had T cell clusters of smaller size and overall, the number of T cells correlated positively with the VI dynamics (**Figure 2**A). Because every mouse received a separate preparation of activated T cells we speculate that the T cells transferred into mouse 3 may have been sub-optimally activated. We know that naive T cells migrate poorly to the liver [41], and it is possible that sub-optimally activated T cells may be less able to locate and eliminate the parasites.

Our estimates of the T cell killing rates were based on the DDR model predicting kinetics of T cell clustering around individual liver stages (**Supplemental Figure S7**). The typical prediction of this model is that the rate of T cell exit from the cluster is larger than the rate of T cell recruitment into the cluster, *μ > λ*_1_. This contradicts the basic idea of the DDR model suggesting amplification of the number of T cells in a cluster via positive feedback loop [17]. We suggested that the parameters of the DDR model are likely to be time-dependent, and with *λ*_1_ *> μ*, the recruitment parameters decline with time since T cell transfer allowing both an increase in number of T cells in larger clusters but also stabilizing T cell numbers at later time points [23]. We simulated T cell clustering dynamics with the DDR-alpha model (DDR model with time-dependent parameters, eqn. (10)) and then fitted the DDR model with time-independent parameters (eqn. (11)) to a subset of these “simulated data”. We found that the DDR model typically overestimates the number of T cells present near the parasite at early time points suggesting that we may have slightly underestimated the T cell killing efficacy (results not shown). Interestingly, a recent study analyzed recruitment of individual CD8+ T cells to tumor-containing organoids and found their data on the T cell cluster dynamics to be best consistent with the DDR model [33].

Given that we do not observe the kinetics of formation of T cell clusters from the first T cell finding the parasite, one could wonder why we need to use the DDR model to estimate T cell killing efficacy. We repeated our analyses (by fitting eqn. (12) with *n* = 1 to VI data) assuming that killing starts only at the time of imaging and used the actual measured T cell numbers per parasite as an input *T* (*t*) into the model by interpolation in Mathematica. Interestingly, we found that for many parasites the estimated killing efficacy is nearly identical to that found by using the DDR model. However, for a fraction (10-15%) we found higher killing rates (results not shown). This is perhaps unsurprising – by limiting the time of interaction between T cells and the parasite to only the observed time period we are likely to overestimate T cell killing efficacy.

Several in vitro studies have suggested that CD8+ T cells are able to serially kill their targets [42–45]. Because we typically followed individual parasites located in different areas of the liver we could not document if T cells in these experiments performed serial killing. In fact, given seemingly random search of most T cells for the liver stages [46], the sparsity of infected hepatocytes in the liver even at high infection doses, and relatively long time it takes for T cells to kill the parasite, it is unlikely that serial killing is relevant to CD8+ T cell-mediated elimination of Plasmodium liver stages in mice.

Our study opens up directions for future research. Details of T cell clustering around individual liver stages remain incompletely understood. While previously we found a strong support for the DDR model [23] our inability to find evidence of attraction of most T cells towards the infection site is troubling [46]. One recent study also suggested a potential role of recirculating T cells in the control of infection with Plasmodium sporozoites [47], and T cells may be following attraction cues via the liver sinusoids rather than via direct, Euclidean distance-based search as we have assumed previously [46]. Developing rigorous frameworks of how T cells search for the infection via the sinusoids is an active area of our research. Several cytokines such as IFNg and chemokine receptors (e.g., CXCR3, CCR5, and others) have been shown to be important in T cell-mediated control of Plasmodium infection [12, 48]. The role of these molecules in the efficiency of T cell search for and elimination of Plasmodium liver stages remains to be investigated. Future studies will need to combine the estimates of the time it takes for CD8+ T cells to locate and then to kill the liver stages. While several studies have found a bias in T cell movement towards an infection site [49, 50], we failed to do so for most T cells in the liver even using a more sensitive metric to detect attraction [46]. In the absence of attracting cues the time to find the infection site may be substantial [12, 46] further reducing the probability that a single T cell is able to eliminate a liver stage and so will require even larger T cell clusters for efficient elimination of the liver stage in 48 hours of its lifespan.

In our current analyses we could not determine if lack of VI decline for some parasites in the presence of sizable CD8+ T cell clusters is due to resistance of the parasite to killing, inability of T cells to recognize the infection, or because clustered T cells are nonlytic. We believe that the first two explanations are unlikely. A large enough population of Plasmodium-specific CD8+ T cells induced by vaccination or adoptive cell transfer is able to provide sterilizing protection against exposure to sporozoites suggesting that no sporozoite is truly resistant to elimination [5, 8, 10]. Clustering of CD8+ T cells is likely to require recognition of the parasite and TCR-mediated signaling since T cells of irrelevant specificity are unable to cluster around Plasmodium liver stages [17, 23]. It is expected that not all activated CD8+ T cells will be cytolytic, and estimating the proportion of T cells that may be nonlytic by measuring expression of surface markers is an important area of future research.

Taken together, our analyses indicate that the average ability of CD8+ T cells to kill Plasmodium liver stages is rather limited and is not in agreement with the conclusion that activated CD8+ T cell act as “rapid” and “serial” killers that destroy hundreds or thousands of targets per day [38]. Deeper understanding of factors that contribute to the ability of T cells to kill their targets – such as the dynamics of CD8+ T cell responses, the cytolytic heterogeneity of CD8+ T cells, T cell recirculation/migration kinetics, and specific details of the tissues where T cells encounter their targets [51] – are likely to improve our ability to estimate T cell kill rates and to make more accurate predictions on the number of T cells required to control infections and cancers [52, 53]. Ultimately, better experimental methods, for example those using Ca-flux mice allowing to visualize recognition of antigen by T cells (and thus potentially, the delivery of the lethal hit) [20], will provide more a sophisticated description of how CD8+ T cells kill their targets in vivo.

## Data availability

The data used in the analysis are provided as a supplement to the paper as well as on github along with computer codes: https://github.com/soumenbera89/Killing-of-Malaria-Parasite-by-CD8-T-cell

## Code availability

We performed most of our analyses in python and provided several key codes to ensure reproducibility of our results on github: https://github.com/soumenbera89/Killing-of-Malaria-Parasite-by-CD8-T-cell

## Author’s contributions

The data for the analyses have been generated in previous study primarily by IAC and RA. RA also contributed previously unpublished changes in the vitality index of Py liver stages over time in control experiments. VVG and SB designed some of the quantitative analyses including formulation of mathematical models. Different analyses have been performed by SB and VVG. IAC provided feedback on the analyses performed. SB wrote the first draft of the paper which was then finalized by VVG and edited by all authors. IAC proofread the paper to ensure proper English. All authors contributed to writing the final version of the paper.

### Abbreviations

Py: Plasmodium yoelii
GFP: green fluorescent protein
VI: vitality index
DDR: density-dependent recruitment

## Acknowledgements

This work was supported mainly by R01GM118553 and partially by R01AI158963 NIH awards to VVG.

## Supplemental Information

### Additional mathematical modeling-based analyses

#### Dose response models to predict parasite’s VI from T cell numbers

In our data both parasite’s VI and T cell number per parasite change over time. In addition to modeling the dynamics of VI change (see below) we aggregated the data and calculated the relationship between the VI and T cell numbers per parasite *T* (*t*) at a given time point *t*. For the analysis we normalized the VI for each parasite to its initial value found at the beginning of imaging. To describe the “dose” relationship between the VI and the number of CD8+ T cells we used several alternative models. These include a log-logistics model:

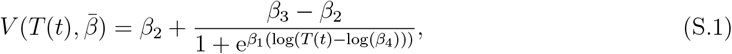

Where 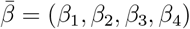 is the vector of parameters, *β*_1_ is the steepness of the dose-response curve, *β*_2_, *β*_3_ are the lower and upper limit of the response, and *β*_4_ is the effective dose *T*_*IC*50_. Other models are

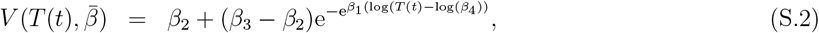

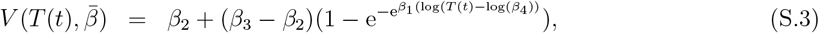

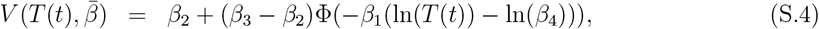

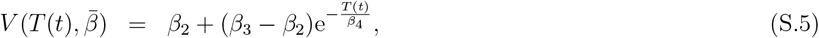

where eqns. (S.2)–(S.5) are for Weibull I, Weibull II, log-normal, and exponential functional response, respectively; Φ is cumulative normal distribution function. We fitted these models to data using R package *drc*. In *drc*, parameters are estimated using maximum likelihood that is under the assumption of normally distributed response simplifies to nonlinear least squares [54]. From the parameter estimates we calculated *T*_*IC*50_ as number of CD8+ T cells required for 50% inhibition of parasite’s VI from its initial value. Typically, *T*_*IC*50_ was computed numerically by setting *V* = 0.5 in the model and solving for *T* (*t*) with estimated parameters 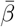.

#### A forecasting statistical model to estimate next VI from previous VI and T cell numbers

To investigate whether CD8+ T cells may impact dynamics of parasite’s VI in subtle ways we borrowed a forecasting approach from economics and several alternative mathematical models from disease population ecology [55]. The basic idea of the model is to use information on the parasite’s VI (*V*_*t*_) and T cell number around the parasite (*T*_*t*_) at time *t* to predict parasite’s VI (*V*_*t*+1_) at the next time instance *t* + 1. We consider the following model:

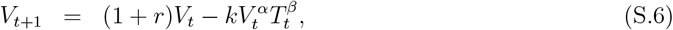

where *r* is the growth rate of the VI and *k* is the reduction rate of the VI due to T cells. Departures from the mass-action killing of the parasite by T cells are modelled with parameter *β* with *β >* 1 indicating cooperation at killing and *β <* 1 indicating competition at killing. We also tested if the model with no T cell-mediated killing (*k* = 0) describes the data with similar quality as the full model.

We fitted these model to the parasite’s VI using linear regression (least squares) in R by taking *V*_*t*_ and *T*_*t*_ given in the data and predicting *V*_*t*+1_ also found in the data. Specifically, for *V*_*t*+1_ we took observed values for VI for a given parasite starting at time point *t* = 2 and regressed this vector against VI values starting at *t* = 1 until *t* = *N −* 1 where *N* is the number of data points for each specific parasite. To investigate if different dynamics of T cell numbers per parasite may allow for a better description of VI dynamics we allowed the number of T cells per parasite i) to change randomly, ii) to increase over time from the lowest number found in the data for a given parasite to 10 (maximal value in our data), and iii) to decrease over time from the highest number of T cells found in the data for a given parasite to zero (lowest value found in our data).

#### A forecasting model with increasing T cell number per parasite better describes change in parasite’s VI

Parasite VI fluctuated over time for most parasites (e.g., **Figure 1Bii**). To investigate whether current VI and T cell number per parasite are predictive of the VI at a later time instance we used a “forecasting model” approach borrowed from economics (**Supplemental Figure S11Ai**). We assumed how current and past VI are related, i.e., whether the next step VI is directly determined by the previous VI value (eqn. (S.6)). In the model we then considered the impact of CD8+ T cell number at the current time moment on the next time VI (time intervals between sequential measurements were between 1.16 to 5.06 min), and examined whether changes of T cell number per parasite improves the quality of the model fits of the VI data. We found that for T cell dynamics observed in experiments, adding T cell dynamics to the model fit improved the prediction of VI dynamics for 6 out of 19 parasites (**Supplemental Figure S11Bi**).

We wondered if there may be errors in measuring of how many CD8+ T cells are located near individual parasites. In our experiments the typical imaging depth is *h* = 50 *μ*m, and given that we count T cells located within *r* = 40 *μ*m from the parasite, the relative volume in which we typically count T cells near the deep parasite is *πr*^2^*h/*(4*/*3*πr*^3^) *≈* 47%. This calculation suggests that we may be underestimating the number of T cells located near individual parasites, and it is possible that our determined change in T cell numbers per parasite may also be not fully accurate.

To investigate whether different T cell dynamics may improve prediction of the VI changes over time we simulated T cell dynamics for each parasite in three different ways: we either 1) assumed that T cell number per parasite increases with time since T cell transfer from the minimal value found for a given parasite to 10 (the maximum observed in our data), or 2) randomly chose a T cell number between the minimal and maximal value, or 3) assumed that T cell number declines from the maximal value over time to zero. We found that increasing the number of T cells over time improved the description of the VI dynamics over the null model with no T cells in 12/19 parasites – a double when compared to the model fits when using actual data (*χ*_1_ = 2.64, *p* = 0.10; **Supplemental Figure S11Bii**). This result was mostly independent of the rate at which T cell numbers were assumed to increase over time. In contrast, choosing random T cell numbers or decreasing T cell numbers over time resulted in fewer fits that improved prediction of parasite’s VI over the null model (**Supplemental Figure S11Biii-Biv**). These results suggest that there may be an increase in the number of T cells around individual parasites that is not fully detectable by the current imaging techniques at least for some (e.g., deep) parasites.

**Supplementary Figure S1:**
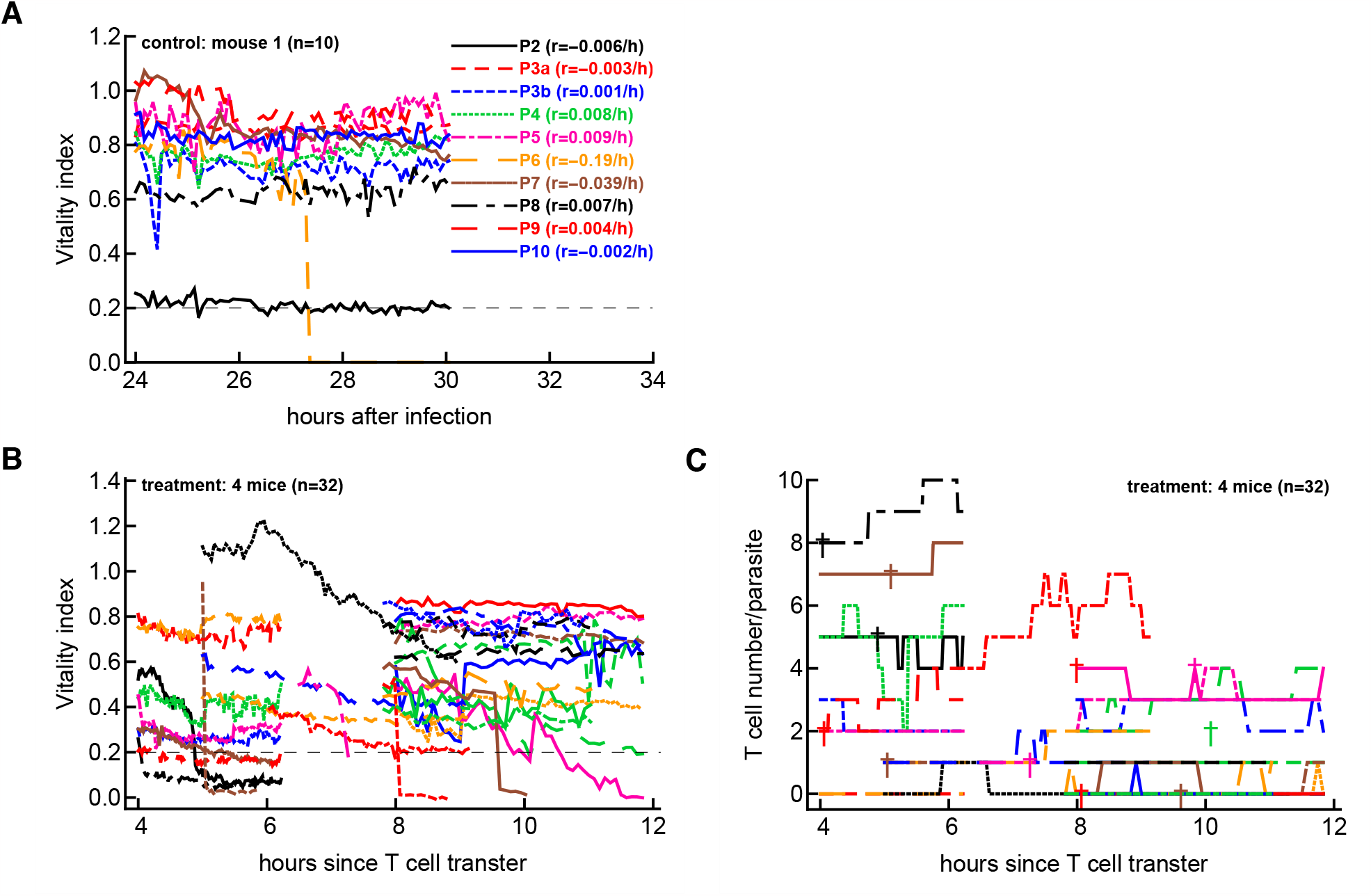
Detailed data on dynamics of VI and T cell numbers in control and treatment groups of mice. (A) We plot the dynamics of parasite’s VI for a subset of parasites (*n* = 10) in one of the two control mice along with estimated rates of VI change over time *r*. (B) We plot the dynamics of parasite’s VI from 32 parasites in 4 mice in treatment group. (C) We plot the dynamics of T cell numbers located near each of the liver stages in B. Crosses indicate parasites that eventually died during the imaging (VI dropped below 0.2). In panel A&B dashed horizontal line denotes the threshold for a dead parasite (VI = 0.2). Note that in panel A time is in hours since infection and in panels B-C time is in hours since T cell transfer (that occurred 20 hours after the infection, see **Figure 1A**). Data for individual parasites are also shown in **Supplemental Figures S2 and S4**.

**Supplementary Table S1:** Estimated parameters of the dose response models. We fitted log-logistics (eqn. (S.1)), Weibull I (eqn. (S.2)), Weibull II (eqn. (S.3)), lognormal (eqn. (S.4)), and exponential decay (eqn. (S.5)) models to normalized VI data divided into different time periods (4-8, 6-8, 8-11, and 4-11 hours post T cell transfer, see Materials and methods for more detail). Model fitting was done using `drc` package in R. Estimated parameters are *β*_1_ (slope), *β*_2_ (lower limit), *β*_3_ (upper limit), and *β*_4_ (*T*_*IC*50_), and NA denotes not applicable. Values for lower and upper limits were restricted to 0 and 100, respectively.

| Model | $\beta_1$ | $\beta_2$ | $\beta_3$ | $\beta_4$ |
| --- | --- | --- | --- | --- |
| 4-11 hours |  |  |  |  |
| Log-logistic | 0.9536 | 0 | 100 | 3.2 |
| Weibull I | 0.7 | 0 | 100 | 5.66 |
| Weibull II | 0.287 | 0 | 100 | 2.386 |
| Lognormal | -0.6 | 0 | 100 | 3.2 |
| Exponential decay | NA | 4.99 | 0 | 100 |
| 4-6 hours |  |  |  |  |
| Log-logistic | 0.47 | 0 | 100 | 0.74 |
| Weibull I | 0.29 | 0 | 100 | 2.39 |
| Weibull II | -0.39 | 0 | 100 | 0.31 |
| Lognormal | -0.29 | 0 | 100 | 0.73 |
| Exponential decay | NA | 2.92 | 0 | 100 |
| 6-8 hours |  |  |  |  |
| Log-logistic | 0.92 | 0 | 100 | 2.02 |
| Weibull I | 0.61 | 0 | 100 | 3.89 |
| Weibull II | -0.7 | 0 | 100 | 1.14 |
| Lognormal | -0.57 | 0 | 100 | 2.03 |
| Exponential decay | NA | 3.27 | 0 | 100 |
| 8-11 hours |  |  |  |  |
| Log-logistic | 15.77 | 0 | 100 | 143.89 |
| Weibull I | 276.2 | 0 | 100 | 7054.2 |
| Weibull II | -2875.7 | 0 | 100 | 7088.4 |
| Lognormal | -749.03 | 0 | 100 | 11301.5 |
| Exponential decay | NA | 18.85 | 0 | 100 |

**Supplementary Figure S2:**
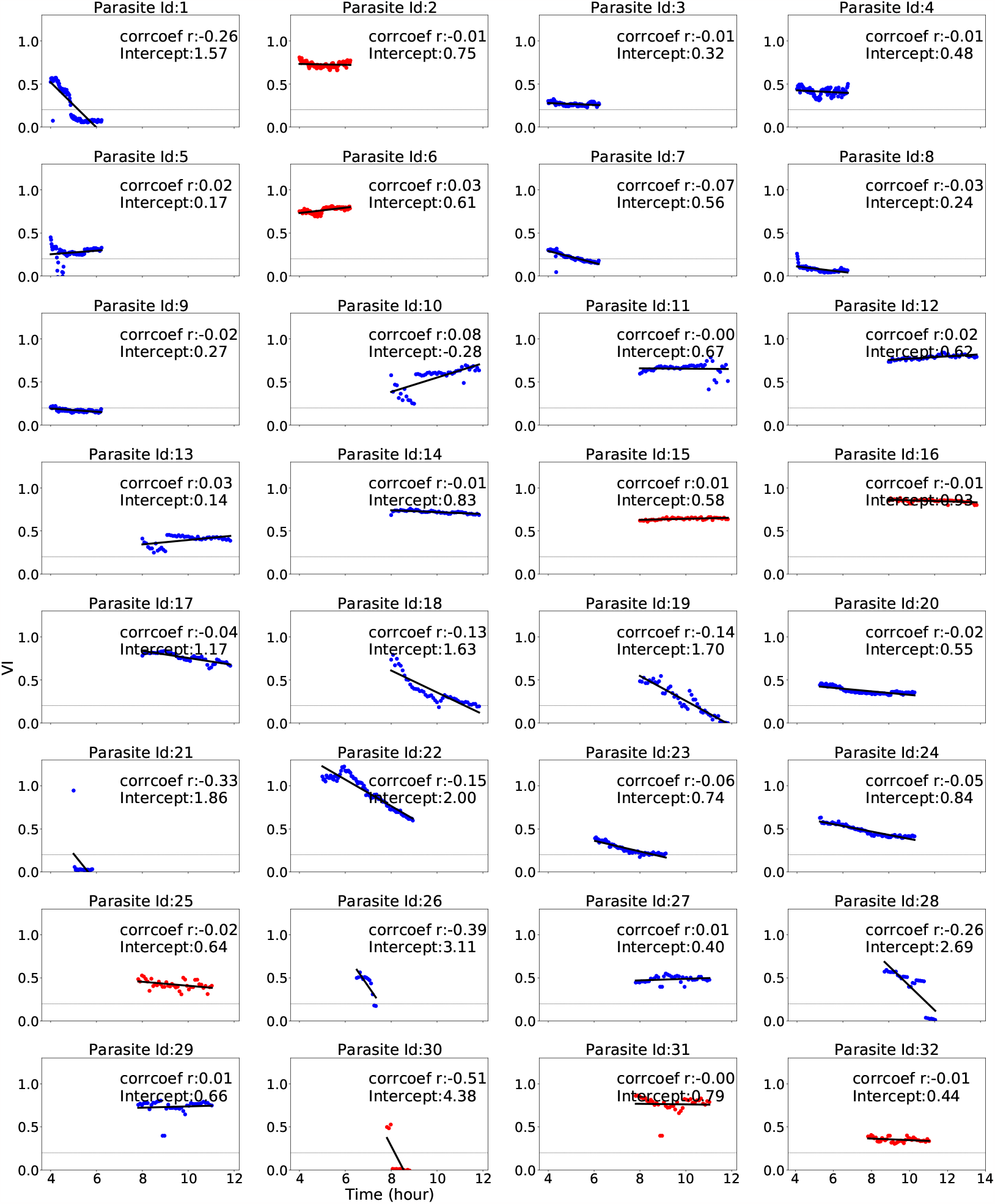
Variable dynamics of the parasite’s VI with time for all 32 parasites in the data. We performed linear regression of VI vs. time and parameters of the regression (correlation coefficient and intercept) are indicated on individual panels. Regressions were performed using **scipy.stat** python package. Dotted horizontal line represent the VI = 0.2 cutoff for dead parasites. Red markers denote time instances for which the number of T cells near the parasite was zero.

**Supplementary Figure S3:**
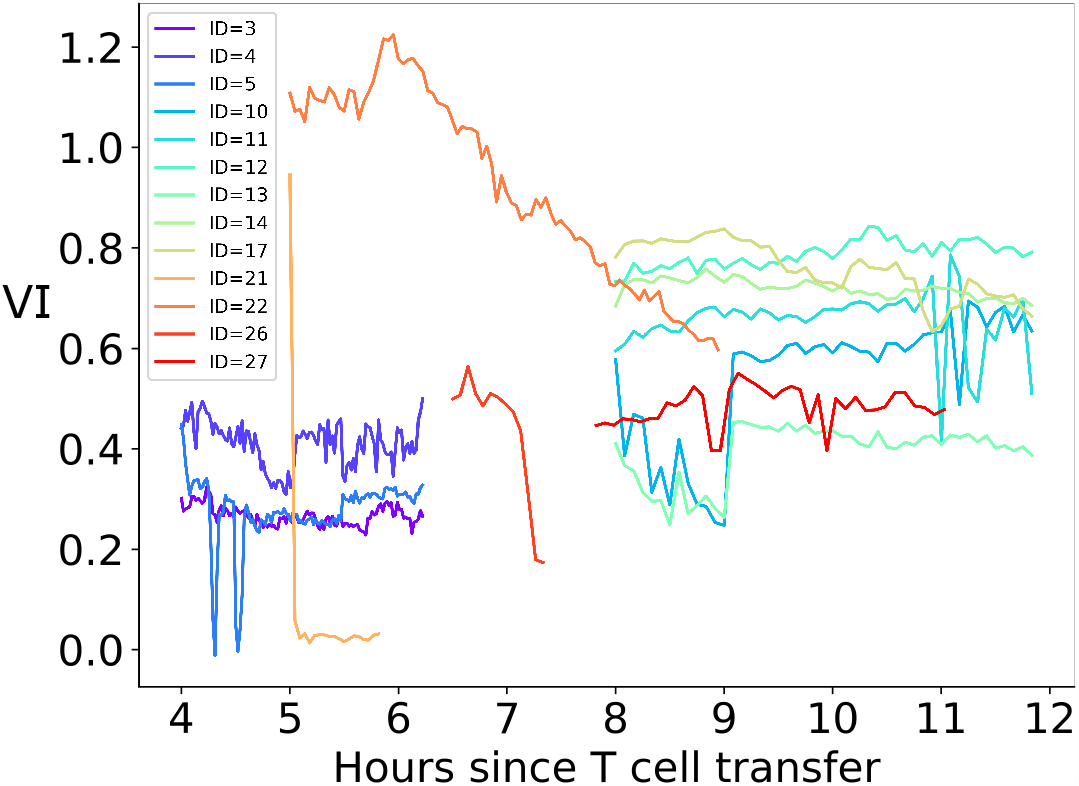
Stability in parasite’s VI dynamics with time for many parasites with non-zero T cells. We show the dynamics of VI of parasites that had some T cells nearby but did not die during the observation period (i.e., VI remained above 0.2). The correlation between VI and T cell numbers for these parasites is non-negative and thus, these were excluded from some of the analyses. The data are colored from violet to red based on the parasite ID number.

**Supplementary Figure S4:**
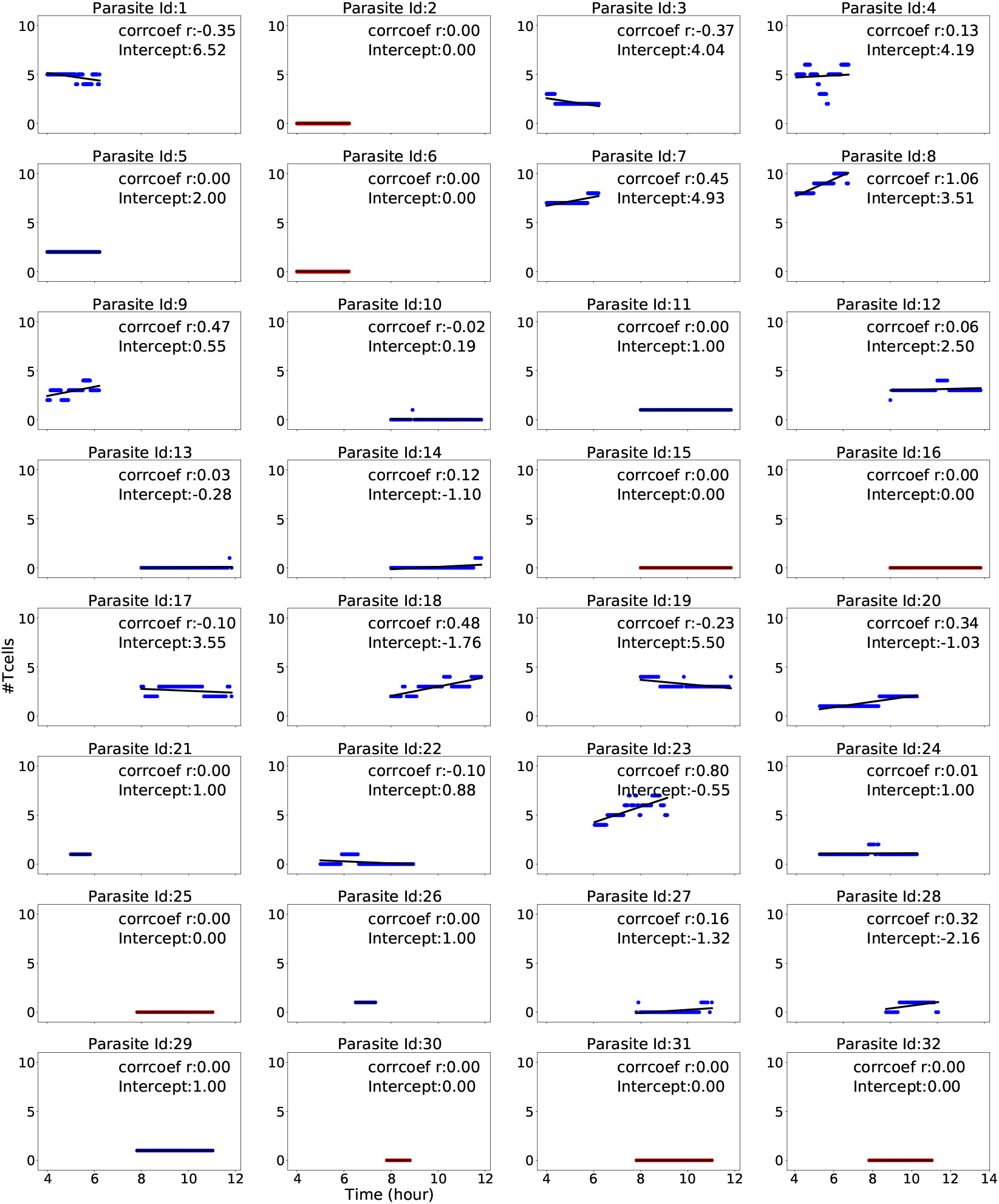
Dynamics of T cells near each of 32 parasites. The correlation coefficient and intercept of the T cell’s trajectory was calculated using *scipy*.*stat* Python package. Dotted horizontal line represent the average number of CD8+ T cells per parasite. Parasites with following IDs did not have any T cells nearby during the imaging period: 2, 6, 15, 16, 25, 30, 31, 32 (denoted by red markers).

**Supplementary Figure S5:**
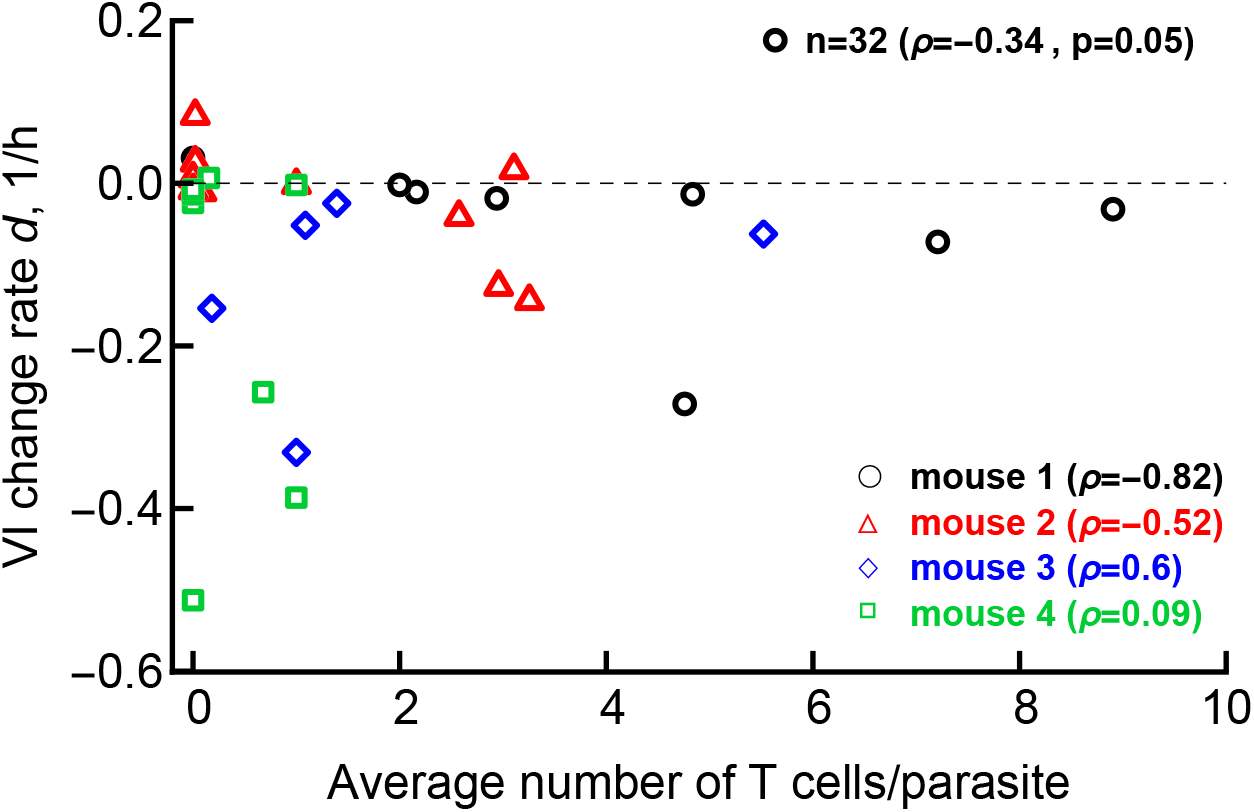
The number of T cells per parasite is a relatively poor predictor of the VI change rate. For every of 32 parasites in our datasets we calculated the rate *d* of VI change with time using linear regression (see **Supplemental Figure S2**) and for the same parasites we calculated the average number of T cells observed in imaging experiments. Negative values for the change rate imply declining VI. Note that many parasites had only few T cells in their vicinity. Spearman rank correlation coefficient *ρ* and p value from the test is shown.

**Supplementary Figure S6:**
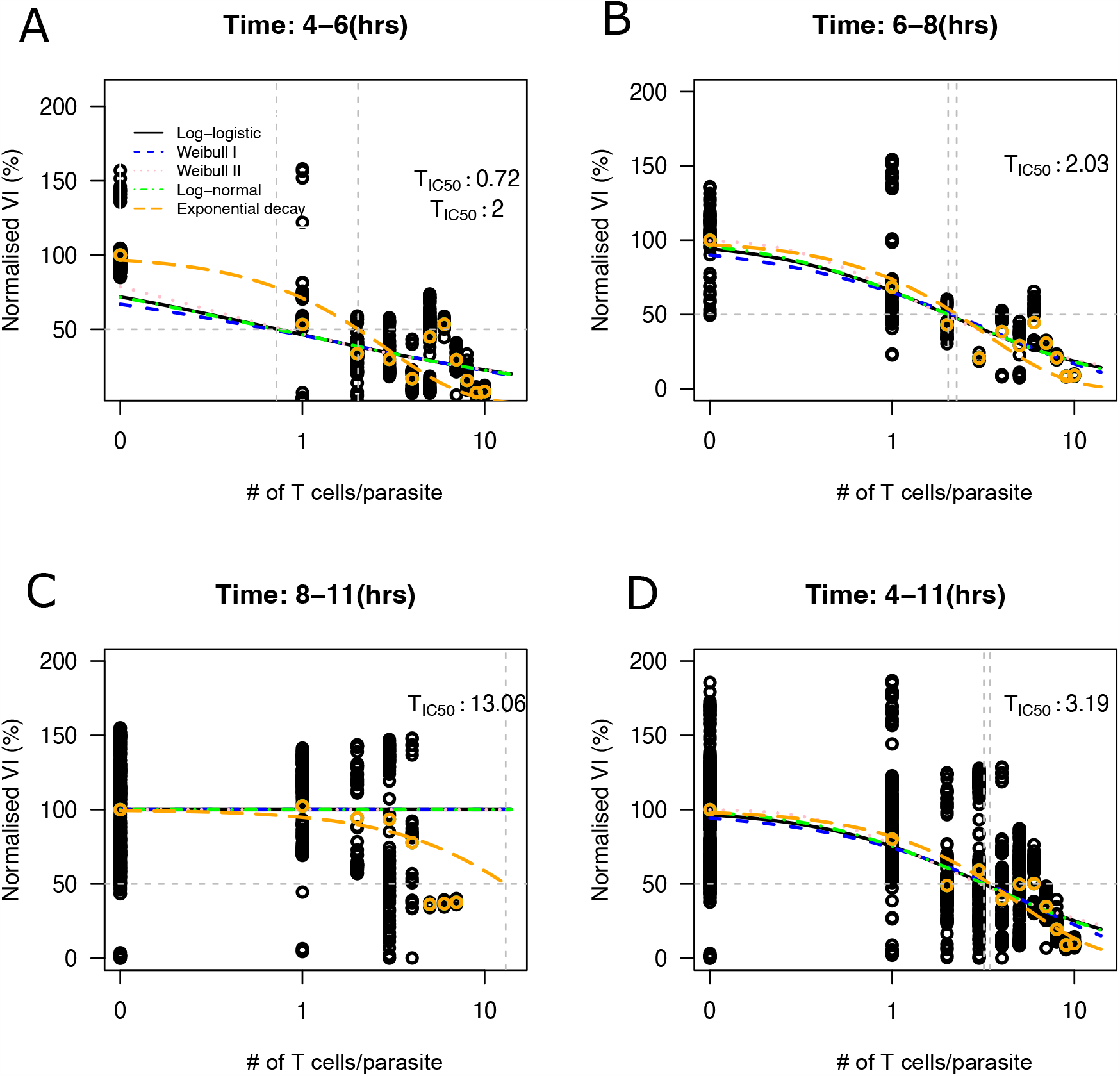
Higher numbers of T cells per parasite are needed to reduce parasite’s VI by 50% at later times since T cell transfer. We fitted several dose-response functions (eqns. (S.1)–(S.5)) to normalized experimental data on VI values for all time points with T cell numbers as a “dose”. Data were normalized to their initial value. (A-C) Datasets were partitioned in different time periods (4-6, 6-8, and 8-11 hours after T cell transfer) (D) The models were fitted to all data. A decision-tree based approaches is used to select the best fit model among the candidates using AIC in R package *drc*. The predicted number of T cells at which VI reaches 50% of its initial value (*T*_*IC*50_) is shown on individual panels by the dashed vertical lines. Dashed horizontal lines denote 50% of initial VI. Parameters of the best fit models are given in **Supplemental Table S1**.

**Supplementary Table S2:** Estimated parameters of the forecasting model in which the number of T cells per parasite was assumed to increase over time at different time intervals. The null model (without T cell, eqn. (S.6) with *k* = 0), or forecasting model with experimentally measured T cell numbers or when we assumed a steady increase in T cell numbers per parasite (eqn. (S.6)) were fitted to data. All the forecasting models were compared with the null model using ANOVA. Model fitting was done by R package *minpack*.*lm*. Statistical significance code shown with bold where ‘***p’*<* 0.001, ‘**p’ *<* 0.01, and ‘*p’ *<* 0.05. *†* represent model fitted with data without convergence.

| Parasite ID | Null Model (no T cell) | Model (with experimental T cells) |  |  | Model (increasing T cell/15 mins) |  |  | Model (increasing T cell/30 mins) |  |  | Model (increasing T cell/60 mins) |  |  |
| --- | --- | --- | --- | --- | --- | --- | --- | --- | --- | --- | --- | --- | --- |
| | P, 1/hr | P, 1/hr | $\alpha$ | k, 1/hr | P, 1/hr | $\alpha$ | k, 1/hr | P, 1/hr | $\alpha$ | k, 1/hr | P, 1/hr | $\alpha$ | k, 1/hr |
| 1 | -0.036 | 0.0877 | 1.555 | 0.0381 | -0.011 | 9.332 | 1.178 | 0.2443 | **1.4139\*\*\*** | 0.0728 | 0.9628 | **1.132\*\*\*** | 0.2206 |
| 3 | -0.0019 | 0.0105 | 9.069 | 177.7 | 0.0196 | **12.07\*\*** | 5876 | 0.0224 | **9.616\*\*** | 389.8 | 0.0369 | **6.55\*** | 15.22 |
| 4 | -0.0015 | 0.0355 | 7.356 | 1.664 | **0.0649\*** | **7.256\*\*** | 1.829 | **0.055\*** | **7.154\*** | 1.633 | 0.06 | 6.136 | 0.888 |
| 5 | -0.0149 | -0.179 | -0.073 | -0.021 | **-0.096\*\*** | **-0.358\*\*** | **-0.0025\*** | **-0.1167\*\*** | **-0.274\*** | **-0.005\*** | **-0.127\*\*** | -0.257 | -0.008 |
| 7 | -0.016 | 0.3475 | **1.334\*\*\*** | 0.083 | 0.3224 | **1.59\*\*\*** | 0.095 | 0.3628 | **1.638\*\*\*** | 0.121 | 0.362 | **1.6381\*\*\*** | 0.121 |
| 8 | **-0.049\*\*\*** | 0.2662 | **1.4627\*\*\*** | **0.1055\*\*\*** | 0.2499 | **1.548\*\*\*** | **0.114\*\*\*** | 0.3052 | **1.4583\*\*\*** | **0.1149\*\*\*** | **0.3052\*** | **1.4583\*\*\*** | **0.1149\*\*\*** |
| 9 | -0.0036 | **0.0521\*\*** | 4.152 | 4.63 | **0.0828\*\*\*** | **7.201\*\*\*** | 882.1 | **0.078\*\*** | **5.529\*\*\*** | 66.15 | **0.072\*** | **4.283\*\*\*** | 9.265 |
| 12 | 0.0013 | 0.008 | 21.14 | 0.2239 | 0.0073 | 27.64 | 0.26 | 0.007 | 29.04 | 0.556 | 0.007 | 28.22 | 0.7046 |
| 13 | -0.005 | -0.0051 | 2.25 | 0.0118 | 0.006 | 23.15 | 2.6e5 | 0.007 | 22.44 | 3.3e5 | 0.007 | 22.44 | 3.3e5 |
| 14 | -9e-05 | 0.0012 | **145.1\*\*\*** | 4.5e20 | **0.0125\*\*** | **30.88\*\*\*** | 29.88 | **0.0105\*** | **33.91\*\*\*** | 187.4 | **0.0076\*** | **31.82\*\*** | 182 |
| 17 | -0.0035 | 0.0028 | 3.0855 | 0.0042 | 0.0279 | **5.4811\*** | 0.0138 | **0.0421\*** | **6.1223\*\*\*** | **0.0316\*** | 0.0457 | **4.7322\*\*\*** | 0.032 |
| 18 | -0.0374 | 0.0419 | **1.8391\*\*** | 0.0586 | **0.172\*** | **2.621\*\*\*** | **0.1624\*\*** | 0.0744 | **2.4047\*\*\*** | 0.0989 | **-0.0265\*** | 63.93 | 150000 |
| 19 | -0.0393 | **-0.4337\*** | **0.7558\*\*\*** | -0.0906 | 0.1968 | **1.9948\*\*\*** | **0.0868\*** | **0.3357\*** | **1.7309\*\*\*** | **0.1374\*** | 0.3447 | **1.466\*\*\*** | 0.1309 |
| 20 | -0.0031 | 0.0089 | 6.6652 | 2.1162 | **0.0351\*\*** | **8.7531\*\*\*** | 13.704 | **0.0259\*** | **9.1719\*\*\*** | 24.67 | 0.0202 | **7.6859\*\*\*** | 7.643 |
| 22 | **-0.0059\*** | -0.0045 | 10.14 | 0.0014 | 0.0074 | **3.068\*\*** | **0.0032\*** | 0.0058 | **3.55\*\*** | **0.006\*** | 0.0197 | **2.717\*\*\*** | **0.015\*\*** |
| 23 | -0.0126 | **0.203\*** | **1.9964\*\*\*** | **0.1476\*\*** | **0.3146\*\*\*** | **2.5811\*\*\*** | **0.353\*\*\*** | **0.3365\*\*\*** | **2.2805\*\*\*** | **0.3202\*\*\*** | 0.1764 | **1.8378\*\*\*** | **0.1182\*** |
| 24<sup>†</sup> | **-0.0055\*** | -0.0036 | 339.6 | 2.6E66 | -0.0011 | **28.34\*\*\*** | 1.6e4 | 0.0184 | **7.6052\*\*\*** | 0.8049 | **0.04\*\*** | **5.517\*\*\*** | **0.5883\*\*\*** |
| 27 | -0.0005 | 0.0023 | 7.2209 | 1.803 | **0.037\*** | **13.97\*** | 108.08 | 0.0304 | **13.43\*** | 153.32 | 0.0125 | 29.24 | 1.5e07 |
| 28 | -0.0455 | -0.0105 | 0.6655 | 0.0457 | 0.0309 | 3.3187 | 0.1683 | 0.0183 | 3.4518 | 0.3911 | -0.0176 | 11.008 | 226.7 |

**Supplementary Figure S7:**
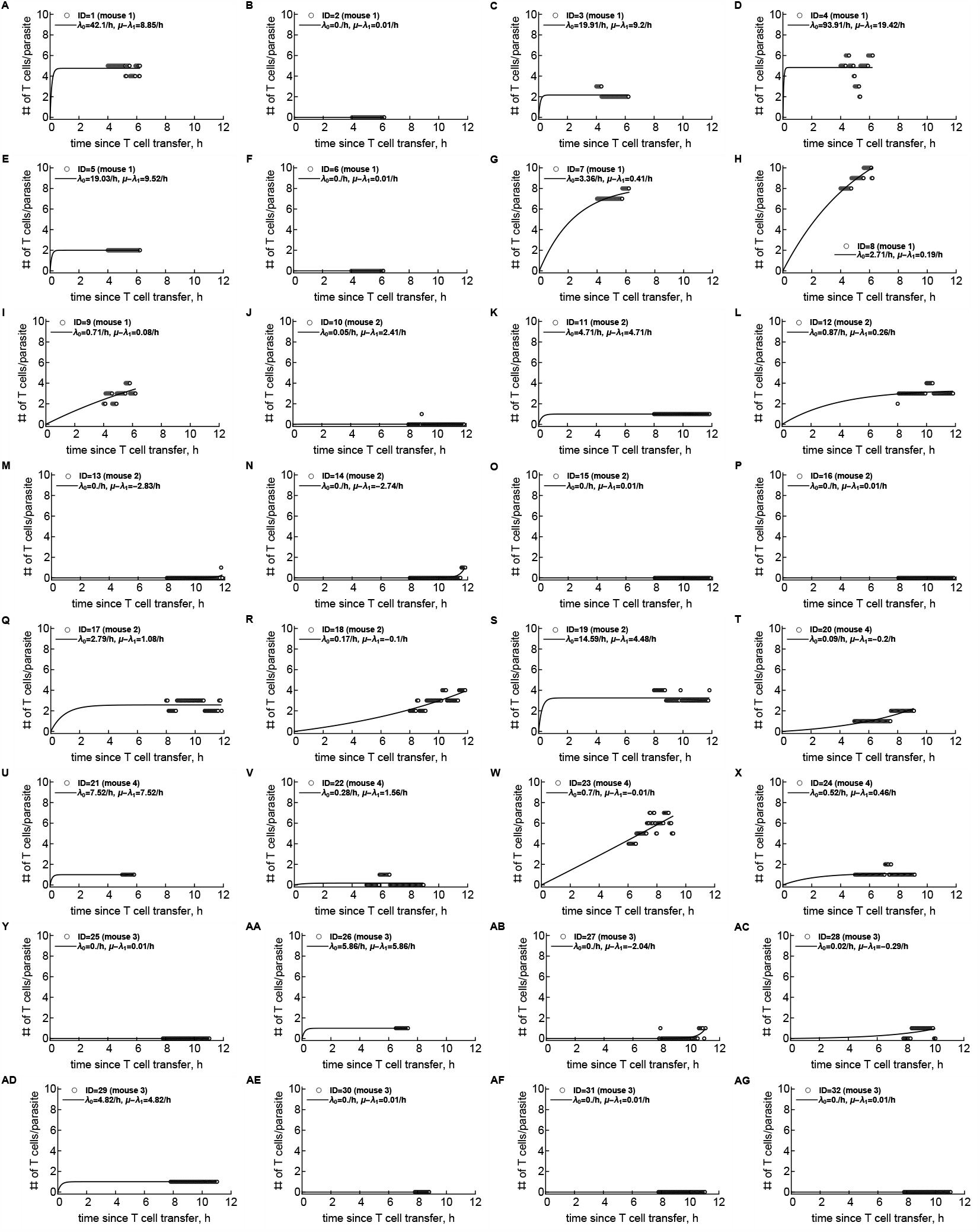
Density-dependent recruitment model describes well the change in the number of T cells different parasites. (A-AG): We fitted the DDR model (eqn. (11)) to the change in T cell number found around individual 32 liver stages. The data are shown by markers and predictions of the best fit model by lines. Each panel shows the estimated parameters *λ*_0_ and *μ − λ*_1_. Fitting was done using `NonlinearModelFit` routine in Mathematica using nonlinear least squares.

**Supplementary Figure S8:**
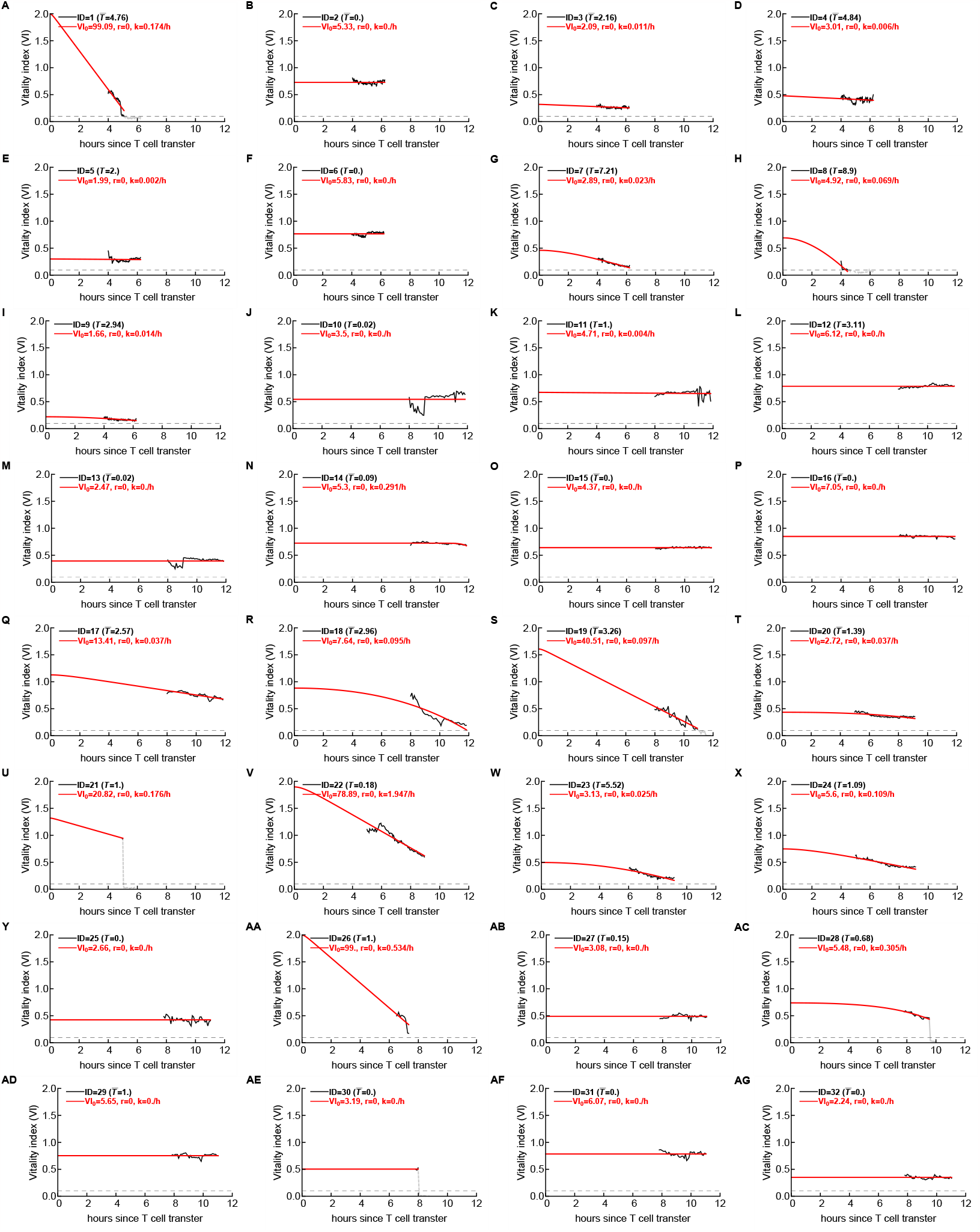
The dynamics of VI is well explained by the mass action-like killing of CD8+ T cells. We fitted alternative models (based on eqn. (12)) to the dynamics of VI for individual liver stages assuming that T cell clustering dynamics following the DDR model (**Supplemental Figure S7)**. (A-AG): The data are shown by black lines and predictions of the best fit basic model (eqn. (12)) by red lines. Each panel shows the time average of the number of T cells per parasite and estimated parameters *V I*_0_, *r*, and *k*. Horizontal dashed line in all panels denotes a critical VI = 0.1 and we only used the data with VI *>* 0.1 in model fitting. A slightly lower critical VI was used here to allow for more data in fitting models to the data. Fitting was done using `FindMinimum` routine in Mathematica with nonlinear least squares. Estimated killing rate is shown up to 3 significant digits; values *k* = 0. are true zeroes.

**Supplementary Figure S9:**
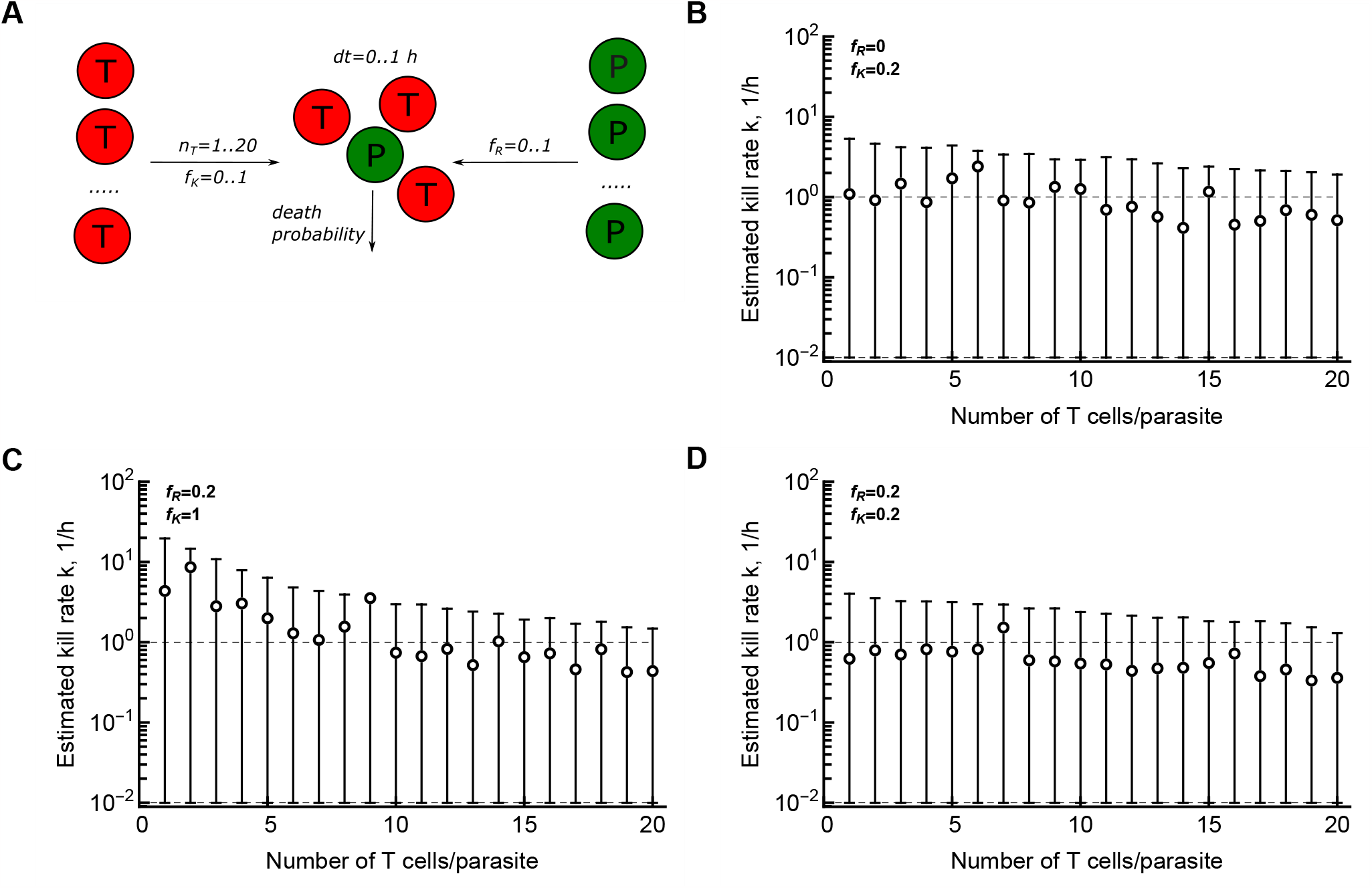
Stochastic simulations suggest that both variability in parasite resistance and per capita T cell killing efficiency explain highly variable estimates of kill rates. (A): Schematic representation of stochastic simulations of killing of the liver stages by T cells. In simulations, we varied the number of T cells in the cluster (*n*_*T*_ = 1 … 20) assuming that T cells either can kill the liver stage at a rate *k* = 1*/*h or are nonlytic *k* = 0 with a probability *f*_*K*_ while the parasite may be fully resistant to killing with a probability *f*_*R*_. T cells interact with the parasite for Δ*t* time chosen uniformly from 0 … 1 h. (B-D): estimated killing efficacy (mean and 95% range) as the function of the number of T cells in the cluster assuming that T cells vary in killing efficacy (*f*_*K*_ = 0.2) while parasites are fully susceptive to killing (*f*_*R*_ = 0, panel B); all T cells can kill (*f*_*K*_ = 1) but some parasites are fully resistant to killing (*f*_*R*_ = 0.2, panel C); both T cells vary in killing efficacy (*f*_*K*_ = 0.2) and parasites vary in resistance (*f*_*R*_ = 0.2, panel D). In panels B-D, in cases when estimated killing efficacy was lower than 10^*−*2^ it was assigned a value 10^*−*2^ to plot these values on the log-scale. Simulations were repeated 10,000 times for each parameter combination.

**Supplementary Figure S10:**
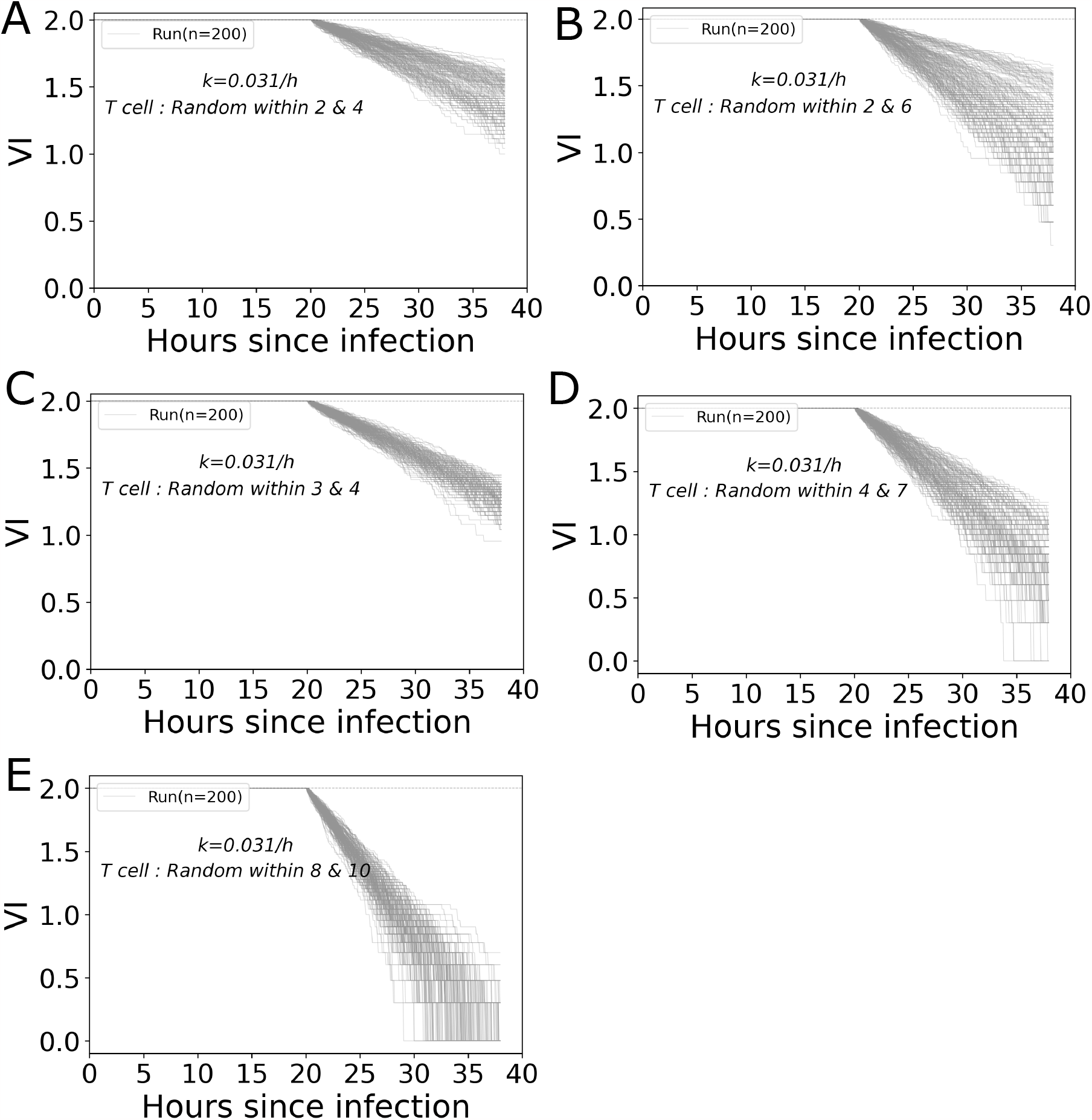
Stochastic simulations of VI dynamics are inconsistent with constant VI observed for most parasites. We performed stochastic simulations of VI dynamics assuming the model with mass-action killing (*k* = 0.031*/*h) using Gillespie algorithm. To mimic our experimental design we simulated the model for 20 hours without any CD8+ T cell with a minimal parasite’s growth rate. Then we adding a random number of T cells (chosen 200 times from a uniform distribution with limits indicated on individual panels and taken from the actual range observed for individual parasites) and simulated the dynamics for the next 16 hours. Dotted line shown the VI threshold of a death parasite. The simulations were done using Python package *gillespy2* with *TauHybrid Solver* routine.

**Supplementary Figure S11:**
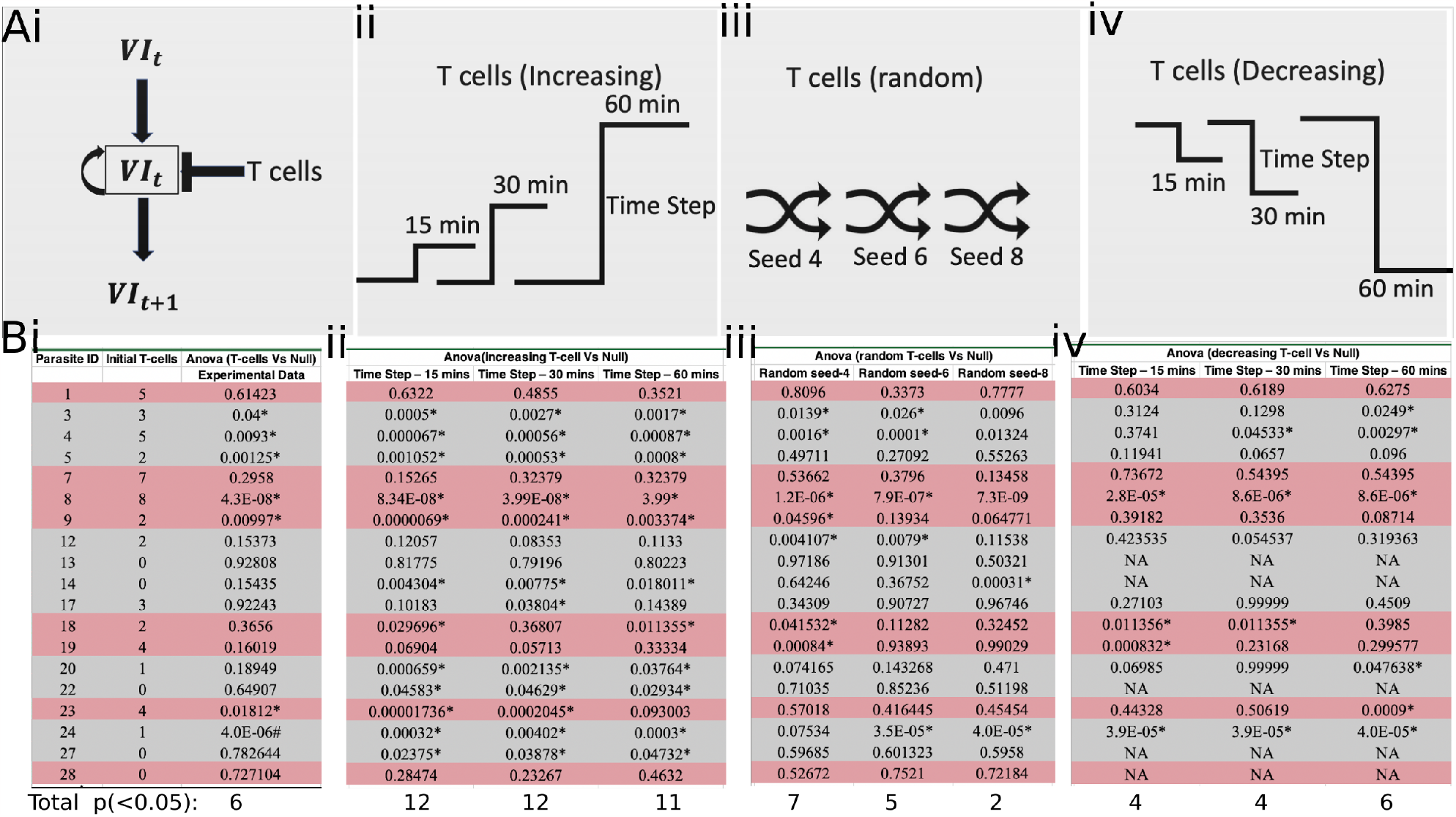
Increasing T cell number in the cluster improves prediction of parasite’s VI over time. (**A**) (i) Schematic diagram of the model to predict next VI (*V*_*t*+1_) using its own previous value (*V*_*t*_) and reduction of VI due to CD8+ T cells present near the parasite. (ii) We increased the T cell number per parasite over time (with three different time steps) starting from the minimal number of T cells found for a given parasite to 10 – a maximum value observed in our data (see **Supplemental Figure S4** for the dynamics of T cell number for each parasite). (iii) We supplied a random T cell number over time (using three different random seeds) that were bounded by lowest and highest T cell number taken from experimental data. (iv) We decreased the T cell number over time (with three different time step) starting from the highest T cell number found in experiments for each parasite to zero. (**B**) We compared the fits of the forecasting null model where the next time step VI is only determined by the VI at the current step (eqn. (S.6) with *k* = 0) with the model in which the dynamics of CD8+ T cell numbers is included (eqn. (S.6)) using ANOVA in R. In fitting the model with changing CD8+ T cell numbers per parasite we (i) took the actual T cell number data for each parasite, (ii) increased CD8+ T cell numbers over time, (iii) chose random T cell numbers over time, and (iv) decreased T cell numbers over time. The analysis was done for 19 parasites for which there was at least 1 T cell nearby at several time points during the imaging period and for which T cell numbers were changing with time. Red and gray shading of rows denotes parasites that are dying (VI has declined or is declining to 0.2) or not dying, respectively. “NA” denotes situation when T cell number was zero. Linear regressions were done in R package *minpack*.*lm* with star *∗* denoting statistically significant difference in model fits at *p <* 0.05. Estimated model parameters are shown in **SupplementalTable S2**.

